# A novel IL-1β reporter assay identifies tolcapone as a caspase-1 suppressor targeting pyroptosis in endotoxemia

**DOI:** 10.64898/2026.07.17.739124

**Authors:** Gabriela Chiritoiu, Simona Ghenea, Gheorghita Isvoranu, Julien Villeneuve, Cristian V.A. Munteanu, Cristian C. Pop, Andreas Bender, Stefana M. Petrescu, Marioara Chiritoiu-Butnaru

## Abstract

Unconventional protein secretion mediated by gasdermin D (GSDMD) pores is essential for the release of pro-inflammatory cytokines such as interleukin-1β (IL-1β), a major driver of many inflammatory pathologies. Despite extensive investigation over the years, discovery of IL-1β secretion modulators has been hindered by the lack of robust, scalable experimental platforms. To date, IL-1β related studies largely rely on primary cells and animal models, suitable for mechanistic studies but not readily scalable for high-throughput applications. Here, we engineered a CRISPR-based reporter cell line that allows quantitative monitoring of endogenous IL-1β secretion while preserving the physiologically relevant inflammasome signaling. This platform faithfully recapitulated the response of primary macrophages to pathogen-associated molecular pattern (PAMPs) stimulation and supported the screening of an FDA-approved drug library comprising 1,398 compounds. Form this screen, we identified tolcapone as a potent inhibitor of IL-1β secretion, reducing cytokine release by more than 80% across the screening pipeline. Mechanistically, tolcapone suppressed caspase-1 activation, thereby limiting GSDMD cleavage, pore formation, and the downstream maturation and secretion of IL-1β and IL-18 *in vitro*. *In vivo*, tolcapone administration attenuated the acute inflammatory response in a lipopolysaccharide-induced endotoxemia model. Together, these findings establish our reporter platform as a robust tool for discovery of endogenous IL-1β secretion modulators and identify tolcapone as a promising inhibitor of inflammasome-driven immune pathology.

## Introduction

Interleukin-1 beta (IL-1β) is a central mediator of innate immunity and major driver in rheumatoid arthritis, atherosclerosis, cancer, hay fever, auto-inflammatory disorders, type 2 diabetes, as well as several neurodegenerative disorders (1,2). Produced primarily by activated macrophages, IL-1β orchestrates inflammatory responses that promote pathogen clearance and tissue repair, but when dysregulated contributes to chronic inflammation and tissue damage (3,4). Although extensively studied, experimental approaches for monitoring IL-1β secretion still rely largely on primary cells and animal models. These systems retain the physiological responses suitable for mechanistic studies, but have limited scalability for systematic discovery of small-molecule modulators of IL-1β release (5,6).

To date, the treatment to control acute and chronic inflammation caused by excessive IL-1β secretion relies on the capacity of recombinant proteins (e.g. Anakinra and Rilonacept) or antibodies (e.g. Canakinumab) to block the binding of extracellular IL-1β to its receptor on target cells to attenuate the propagation of inflammation (5,7). However, the efficiency of these molecules to attenuate IL-1β-induced inflammatory processes in clinical trials did not meet the initial expectations (8,9). An assay that quantitatively tracks endogenous IL-1β secretion while preserving native inflammasome signaling would therefore provide an important platform for both mechanistic studies and drug discovery.

IL-1β, like other members of the IL-1 cytokine family such as IL-1α and IL-18, is synthesised in the cytoplasm of macrophages as precursor molecule (*pro*IL-1β) upon exposure to pathogen-associated molecular patterns (PAMPs) or damage-associated molecular patterns (DAMPs) that activate the TLR (Toll-like receptors)-NF-kB pathway (10,11). A second stimulus, such as release of ATP (adenosine triphosphate) from damaged mitochondria, is required to trigger inflammasome assembly and caspase-1 auto-activation (12–15). This in turn cleaves *pro*IL-1β to produce the bioactive form of the cytokine (*m*IL-1β) which is exported to the extracellular space through the unconventional protein secretion pathway (UPS) (16,17), a tightly regulated process with a prominent role in innate immunity, particularly inflammasome-mediated cytokine release (18,19). Activated caspase-1 also promotes cleavage of gasdermin D (GSDMD), whose N-terminal fragment (GSDMD p30) forms membrane pores that drive cytokine release and mediate an inflammatory form of cell death called pyroptosis (20). GSDMD-dependent pore formation has therefore emerged as a key effector mechanism linking inflammasome activation to the unconventional secretion of IL-1 family cytokines, S100 proteins and other inflammatory mediators (21,22).

To overcome current experimental limitations related to IL-1β secretion, we developed a CRISPR-engineered macrophage reporter cell line by introducing a small tag (HiBiT) into the endogenous IL-1β locus, that allows quantitative monitoring of cytokine secretion. This reporter cell line closely phenocopies the response of primary macrophages to sequential inflammasome stimulation and is suitable for screening applications. Since the compounds can be applied simultaneously with ATP, the assay is particularly well suited to identify molecules that inhibit the export of IL-1β from macrophages rather than upstream cytokine synthesis. Upon screening of an FDA-approved compound library we identified tolcapone as consistently inhibiting IL-1β-mediated inflammatory phenotype reducing cytokine release by more than 80% across the screening pipeline. We further demonstrate that tolcapone impairs caspase-1 activation, limits GSDMD cleavage, and attenuates IL-1β-driven inflammatory responses *in vitro* and *in vivo*. Together, our findings provide a versatile platform for the discovery of endogenous IL-1β secretion modulators and identify tolcapone as a candidate regulator of inflammasome signaling.

## Results

### Generation of a HiBiT-based reporter cell line for monitoring IL-1β secretion

To establish a physiologically relevant system for quantitative monitoring of endogenous IL-1β secretion, we looked for a macrophage cell line that retained the responsiveness to inflammatory stimuli. The commercially available murine macrophage line J774A.1, has been reported to constitutively express and secrete IL-1β, therefore we assessed its capacity to secrete endogenous IL-1β as previously reported (23,24). Following priming with 0.5 μg/ml lipopolysaccharide (LPS) for 16h, J774A.1 cells were stimulated with 2 mM adenosine triphosphate (ATP) and IL-1β secretion was quantified by ELISA at the indicated time points. Endogenous IL-1β was efficiently secreted and became detectable in the culture medium within 30-45 minutes of ATP-stimulation (Fig. 1A).

**Figure 1:**
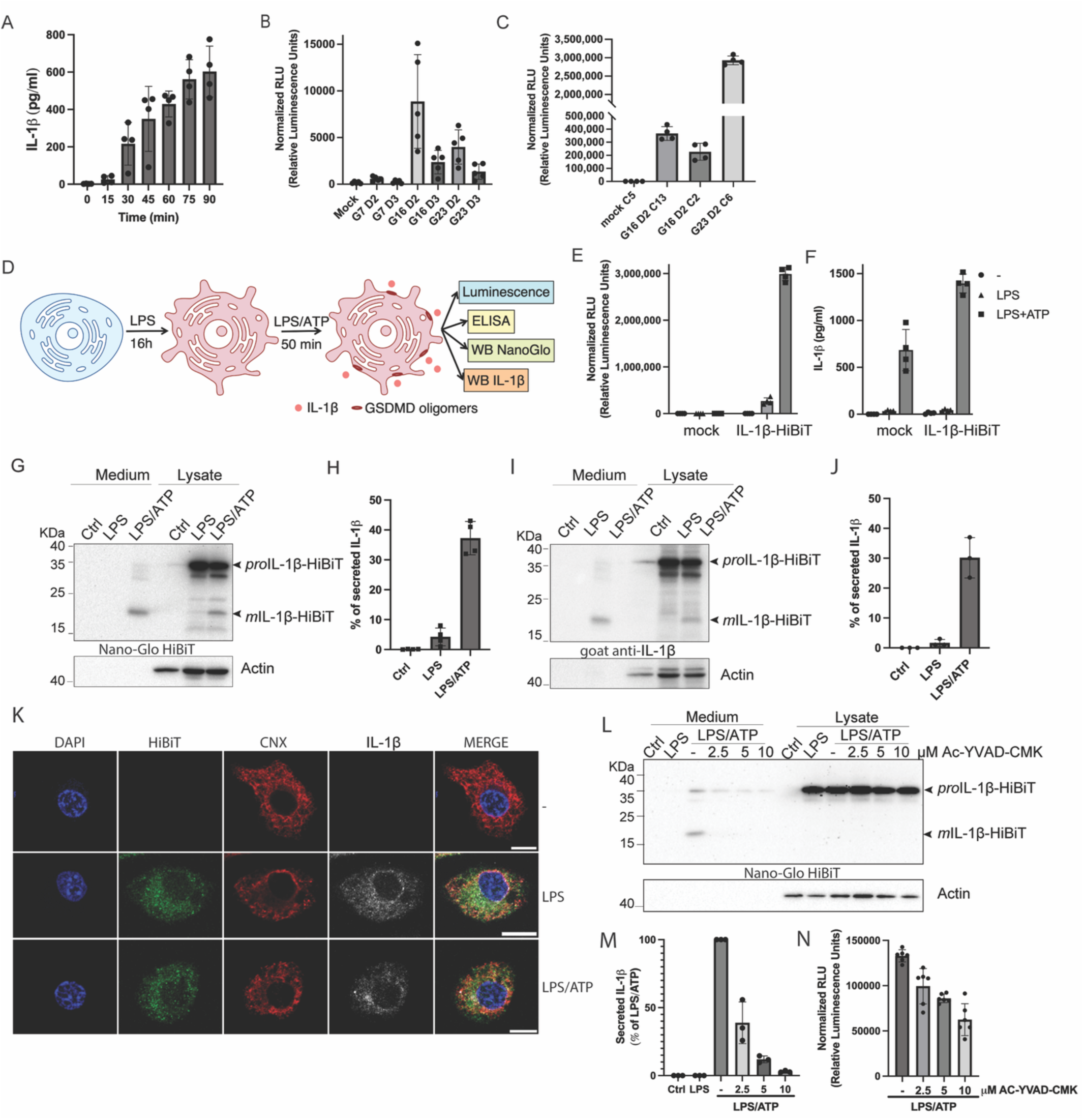
Generation of HiBiT-tagged IL-1β-secreting reporter cell line. **A**. J774.A1 cells were stimulated or not with LPS (0.5 μg/ml) and ATP (2 mM) for the indicated time points. Extracellular medium was collected and processed for sandwich ELISA with IL-1β antibodies according to the manufacturer’s instructions. Values of four independent experiments are represented graphically (*mean ± SD, n=4*); **B.** J774.A1 transfected to express IL-1β-HiBiT (J774.A1-IL-1β-HiBiT) with different combinations of donors and guides, were stimulated with LPS and ATP and the corresponding luminescence to the extracellular medium was detected. Values of five independent experiments are represented graphically (*mean ± SD, n=5*); **C.** Homozygous clones identified by PCR were evaluated for their secretory capacity of IL-1β-HiBiT by luminescence reading. Values of four independent experiments are represented graphically (*mean ± SD, n=4*); **D.** Macrophage stimulation protocol for J774.A1-IL-1β-HiBiT expressing cells; **E, F.** Extracellular medium of cells stimulated as described in (D) were used for luminescence detection (E) or sandwich ELISA using goat anti-IL-1β antibodies (F). Values of four independent experiments are represented graphically (*mean ± SD, n=4*); **G, H.** Lysates and extracellular medium of cells stimulated as described in (D) were separated by SDS-PAGE and transferred on nitrocellulose membrane, followed by detection with the Nano-Glo HiBiT blotting system (G) and band intensity quantification of four independent experiments are presented (H) (*mean ± SD*, n=4); **I, J.** The same samples analysed in (G) were processed for WB and detected with goat anti-IL-1β antibodies (I); rabbit anti-β actin as loading control. Representative images of four independent experiments are presented (J) (*mean ± SD*, n=4 (I) and *mean ± SD*, n=3 (J)); **K.** J774.A1-IL-1β-HiBiT were stimulated as described in (D), fixed with PFA 1% and processed for immunofluorescence using mouse anti-HiBiT antibodies (green), rabbit anti-calnexin antibodies (red), goat anti-IL-1β antibodies (grey), nuclei were stained with DAPI (blue); **L, M.** Cells were seeded and stimulated as described in (D) in the presence or absence of increasing concentrations of the caspase-1 inhibitor (Ac-YVAD-CMK). Extracellular medium and cell lysates were separated by SDS-PAGE, transferred on nitrocellulose and blotted with the Nano-Glo HiBiT blotting system. Actin was used as loading control. Representative images of three independent experiments are presented graphically (*mean ± SD, n=3*); **N.** Medium of cells stimulated as in (L) was used for luminescence activity determination and represented as normalized RLU (relative luminescence units) vs drug concentration. Values of six biological replicates from three independent experiments are represented in (*mean ± SD, n=6*);

Having identified a suitable cell line, we next designed a CRISPR/Cas9-based strategy to engineer J774A.1 cells to express IL-1β fused to a small tag (HiBiT-11 amino acids) minimizing the modifications to the endogenous protein sequence (25,26). The resulting knock-in reporter cell line should produce HiBiT-tagged IL-1β in response to LPS and ATP stimulation, facilitating a rapid quantification of the secreted cytokine in the extracellular medium by complementing HiBiT with extracellular LgBiT to yield luciferase activity (Fig. S1A). J774A.1 cells were electroporated with RNP complexes composed of recombinant Cas9 and one of three guide RNA (Guides: G7, G16, G23) together with single-stranded DNA oligo nucleotide donor (ssODN donors: D2 or D3) for homology-directed repair; a non-targeting crRNA sequence served as mock control (all sequences are described in detail in Table 1). Genome editing was assessed 48h post-transfection by PCR genotyping and by luminescence-based detection of secreted IL-1β-HiBiT respectively (Fig. S1B and Fig. 1B). To minimize variability and avoid selective expansion of unedited cells during passaging, we isolated and expanded single-cell clones. PCR genotyping of 50 clones identified several homozygous HiBiT insertions and the representative results for 12 clones are presented in Fig. S1C. Homozygous clones were evaluated for IL-1β-HiBiT secretion (Fig. 1C) and the clone with the best secretion signal (G23D2 C6-abbreviated IL-1β-HiBiT from here on) was selected for further characterization. For this clone, we first tested the cell number-dependency of the luminescent signal and observed a proportional increase in HiBiT luminescence with increasing cell number, supporting the quantitative performance of the reporter system (Fig. S1D).

**Table 1:**
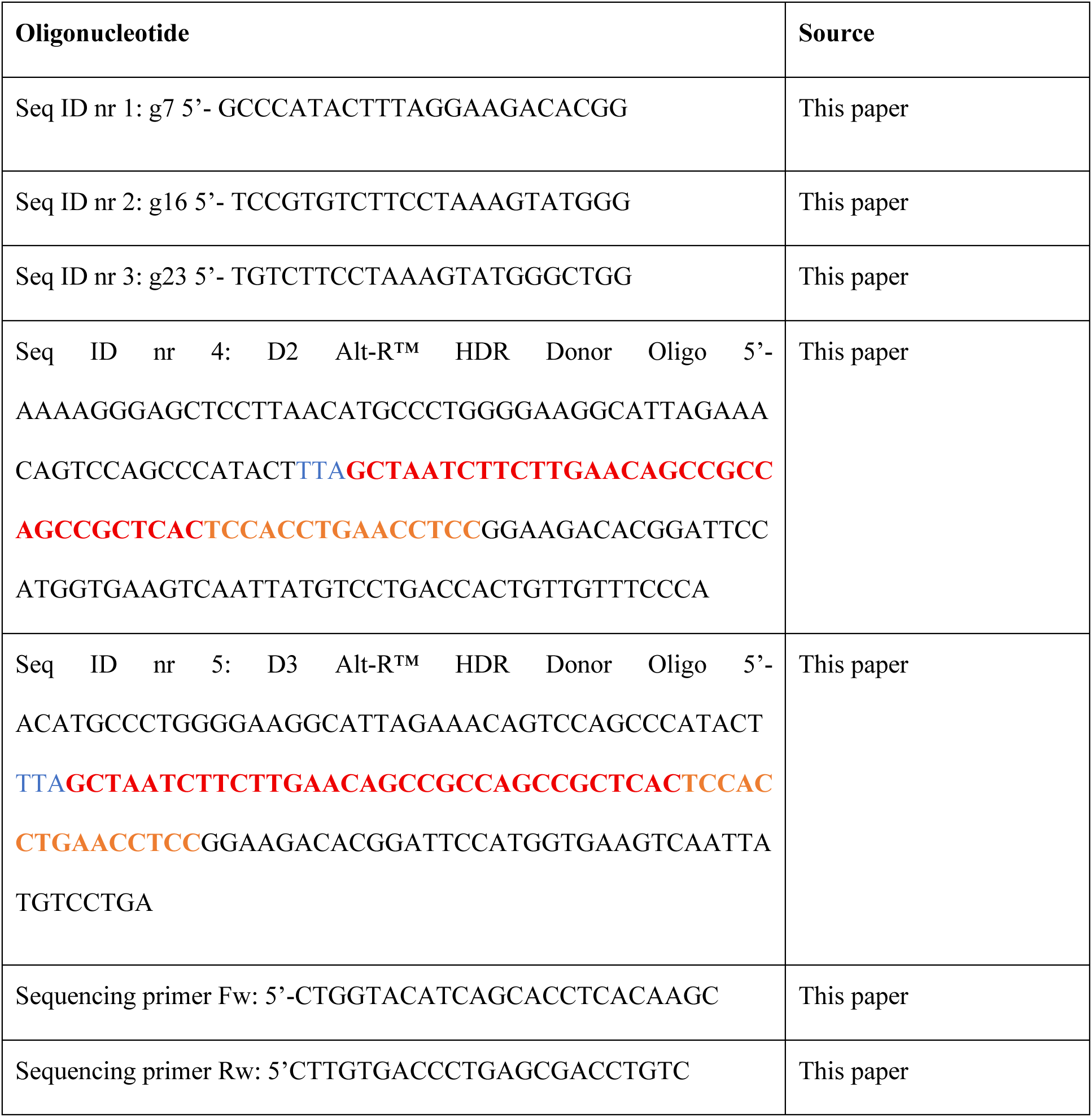
Oligonucleotide sequences.

**Table 2:** Antibodies used in the study.

| Antibodies | Source | Catalog no | WB dilution | IF dilution |
| --- | --- | --- | --- | --- |
| Goat anti-IL-1β | R&D systems | AF-401-NA | 1:1000 | 1:1000 |
| Mouse anti-HiBiT | Promega | N7200 | 1:1000 | 1:800 |
| Phalloidin-AF488 | Invitrogen | A12379 | N/A | 1:200 |
| Rabbit anti-Calnexin | Abcam | ab22595 | 1:5000 | 1:1000 |
| Rabbit anti-Actin | Sigma-Aldrich | A2066 | 1:5000 | N/A |
| Rabbit anti-IL-18 | Proteintech | 10663-1-AP | 1:1000 | N/A |
| Rabbit anti-IL-1α | Proteintech | 16765-1-AP | 1:1000 | N/A |
| Rabbit anti-GSDMD | Abcam | EPR19828 | 1:1000 | N/A |
| Rabbit anti-IL-1β | Proteintech | 16806-1-AP | N/A | 1:1000 |
| Rabbit anti NLRP3 | MedChemExpress | HY-P80246 | 1:1000 | N/A |
| Rabbit anti-ASC/TMS1 | MedChemExpress | HY-P80548 | 1:1000 | N/A |
| Rabbit anti-cleaved caspase-1 | MedChemExpress | HY-P80622 | 1:1000 | N/A |
| Mouse anti-p62/SQSTM1 | Abnova | H00008878-M01 | 1:1000 | 1:800 |
| Rabbit anti-LC3B | Sigma | L7543 | 1:1000 | N/A |

Further, we validated the IL-1β–HiBiT reporter cell line by comparing HiBiT-based and antibody-based detection of IL-1β from the same samples, using the workflow outlined in Fig. 1D. Following stimulation with LPS and ATP to induce IL-1β production and release, culture medium and cell lysates were collected and analysed by ELISA, Western blotting and HiBiT-based detection (luminescence reading and Western blotting detection). Luminescence corresponding to secreted IL-1β-HiBiT was detected exclusively in the culture medium of the reporter cell line and was absent from the mock edited control cell line (Fig. 1E). By contrast, ELISA with IL-1β-targeting antibodies performed on the same samples detected the cytokine in the medium of both mock and edited cell lines (Fig. 1F). We next compared HiBiT-based and antibody-based detection by Western blotting using matched lysate and medium fractions from stimulated cells. IL-1β–HiBiT was detected with comparable specificity both in the extracellular medium and in the cell lysates using either the Nano-Glo HiBiT blotting system (Fig. 1G and H) or the conventional immunoblotting with anti-IL-1β antibodies (Fig. 1I and J). To assess intracellular localisation, we performed immunofluorescence using antibodies against either the HiBiT tag or the C-terminus of IL-1β in stimulated and unstimulated cells. Both the HiBiT and IL-1β-recognizing antibodies revealed a similar intracellular staining pattern after stimulation, whereas no signal was detected in unstimulated cells (Fig. 1K and S1E). Because the secretion of the bioactive IL-1β depends on caspase-1-mediated processing we next asked whether IL-1β-HiBiT was similarly cleaved in the reporter cells (27,28). To test this, stimulated cells were incubated with a selective caspase-1 inhibitor (Ac-YVAD-CMK) which decreased the secretion of IL-1β-HiBiT in a dose dependent manner as assessed by both Western blotting (Fig. 1L and M) and luminescence-based detection (Fig. 1N). Together, these results suggest that the reporter cell line produces and secretes mature IL-1β-HiBiT in response to endogenous inflammasome activation and can be used as an efficient screening system.

### Large-scale screening identifies tolcapone as an inhibitor of IL-1β secretion

#### Optimization of the screening assay

We aimed identify inhibitors that block either the proteolytic processing or IL-1β export across the plasma membrane, therefore, to focus on these last steps the chemicals were added simultaneously with ATP to trigger inflammasome activation following a workflow described in Fig. 2A. First, we optimized cell density and stimulation conditions for cells grown in 96-well plates and measured the luminescence of the reconstituted luciferase (HiBiT+LgBiT) in the supernatant of stimulated cells after 50 minutes. The system was calibrated using DMSO (dimethylsulfoxide, Vehicle control) as negative control and the Caspase-1 inhibitor (AC-YVAD-FMK) as positive control. Compound toxicity was not evaluated separately in this assay, as toxic compounds are expected to induce mitochondrial damage, reactive oxygen species formation, and eventually cell rupture, thereby artificially increasing IL-1β-HiBiT release. Accordingly, compounds associated with hypersecretory phenotypes were excluded from further analysis.

**Figure 2:**
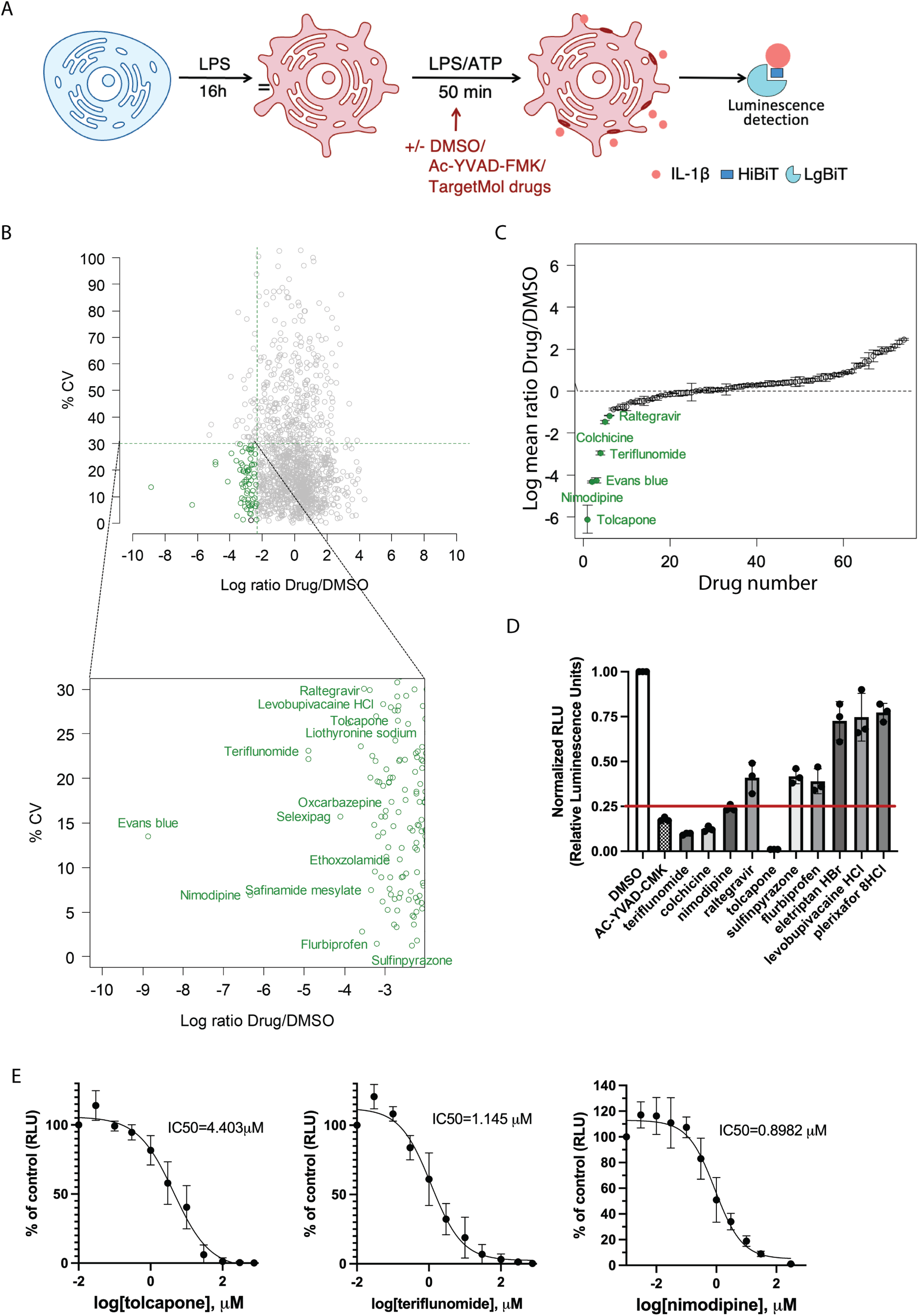
High-throughput screening of FDA approved drugs. **A.** Cell stimulation protocol for high-throughput screening. J774.A1-IL-1β-HiBiT cells were stimulated for 50 min with LPS (0.5 μg/mL) and ATP (2mM) along with DMSO, Ac-YVAD-CMK (10 μM) or TargetMol drugs (20 μM). Fraction of the extracellular medium was used for luminescence measurement using the Nano-Glo HiBiT Extracellular Detection System; **B.** Primary screening results for IL-1β-HiBiT inhibition efficiency (20 μM)). The graph displays CV (%) and log ratio Drug/DMSO. In green are denoted primary hits (log2 ratio of drug/DMSO ≤-2,32 with a coefficient of variation threshold of maximum 30% for triplicates). The inset shows the labelling of the primary hits; **C.** Secondary screening to identify compounds with higher potency and to eliminate the false positive of primary screen preselected 74 compounds. Compounds were selected by considering two cut-offs: log2 ratio of Drug/DMSO ≤-0.4775 to delineate molecules with low potency (30-50% inhibition) and a second cut-off for compounds with log2 ratio of Drug/DMSO≤-1.16, corresponding to molecules with high potency in inhibition (≥55%), labelled in green; **D.** The ten most active compounds selected in the secondary screen were retested at the 10 μM concentration and the normalised luminescence values are represented as bar plots for mean of three independent experiments (*mean ± SD, n=3*); **E.** Dose-response curves for three drugs which showed at least 75% inhibition in the previous experiment (D). The cells were stimulated in the same conditions as for screening only the concentration of the inhibitors were modified with serial dilution from 900 μM to 30 nM inhibitor concentration;

#### Screening of an FDA approved compound library

We chose to screen a commercially available library of FDA-approved drugs with the aim to identify already approved compounds that could be repurposed as anti-inflammatory molecules. Using the optimized assay, we tested 1,398 compounds from the TargetMol library by quantifying luminescence in the supernatants of stimulated cells exposed to each compound at a final concentration of 20 μM (Z’ factor = 0.56 ±0.05). Several of the library compounds that displayed inhibitory capacity were either classical anti-inflammatory drugs or compounds previously reported to affect IL-1β secretion, like flurbiprofen, betamethasone or diclofenac, which further validates the robustness of our assay (29–32). Our primary screen identified 74 compounds that reduced the luminescent signal in the supernatant by more than 80%, (log_2_ ratio of drug/DMSO ≤-2,32) with a coefficient of variation threshold of maximum 30% for triplicates (Fig. 2B and supplementary table 1). These candidates were advanced to a secondary screen at a lower concentration (10 μM) to reduce false positives. In the secondary screen, 11 compounds retained more than 70% inhibitory activity (log_2_ ratio of drug/DMSO≤-0.4775) (Fig. 2C and supplementary table 1). One compound, Evans blue, was excluded because it interfered with the luminescence readout.

The remaining compounds were re-evaluated, and four the most potent compounds showing at least 75% inhibition at 10 μM were selected for further validation (Fig. 2D). These compounds were repurchased from another supplier (MedChem Express) and we reassessed their inhibitory activity using the same experimental set-up. All four drugs consistently inhibited IL-1β-HiBiT secretion with different potencies and teriflunomide and tolcapone exhibited the best dose-dependent effect (Fig. S2A). IC50 values could be determined for 3 compounds (nimodipine, teriflunomide and tolcapone) (Fig. 2E), whereas luminescence measurements for colchicine showed high variability and did not allow reliable IC50 estimation.

#### Validation of assay specificity across secretory cargoes

To further validate the specificity of the HiBiT-based secretion assay, we exploited previously established HiBiT reporter systems in human HeLa cells for both conventionally secreted (PAUF-pancreatic adenocarcinoma up-regulated factor) and unconventionally secreted cargoes (Tau-tubulin associated unit, FGF2 (fibroblast growth factor 2) and human *m*IL-1β (33,34). Treatment with tolcapone for 30, 60 and 120 min did not affect the secretion of PAUF, Tau and FGF2 (Fig. S2 B-D), whereas the secretion of human *m*IL-1β was visibly reduced (Fig. S2 E). These results support the specificity of tolcapone in targeting cytokine release in heterologous systems.

### Validation in primary macrophages confirms tolcapone as lead inhibitor

We next assessed whether the most potent compounds identified in the screening assay retained their inhibitory activity and were able to modulate endogenous IL-1β secretion in primary macrophages using an workflow depicted in Fig. 3A. Tolcapone reduced IL-1β secretion in a dose-dependent manner when added during ATP stimulation, as shown by both Western blotting and ELISA (Fig. 3B-D), thereby validating the results obtained with the reporter cell line. By contrast, nimodipine had a mild but not significant effect on IL-1β secretion in primary macrophages (Fig. S3A), whereas teriflunomide exerted a modest dose-dependent inhibitory effect, as assessed by ELISA and Western blotting (Fig. S3A and B).

**Figure 3:**
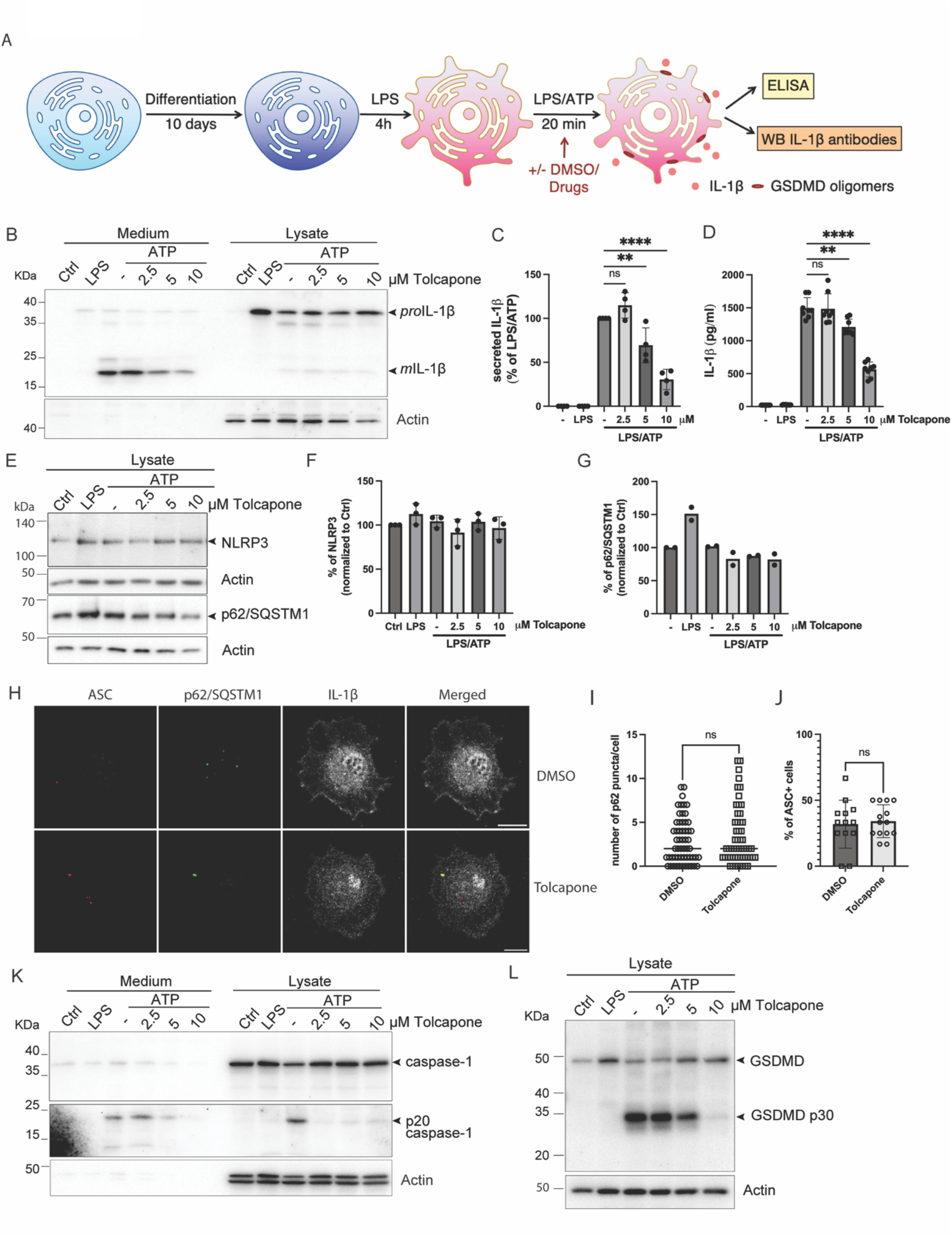
Tolcapone limits GSDMD proteolysis to impair IL-1β secretion. **A.** Protocol for primary macrophage stimulation. Differentiated macrophages were stimulated with LPS (0.5μg/mL) and ATP (2mM) along with DMSO or the indicated compounds. Secreted and intracellular levels of IL-1β and other proteins were evaluated by ELISA or Western blotting; **B.** BMDMs were stimulated as described in (A) in the presence of increasing concentrations of tolcapone. Supernatants and lysates were separated by SDS-PAGE, transferred on nitrocellulose membranes and blotted with antibodies against IL-1β or actin; **C.** Band densitometry of four independent experiments presented in (B) (*mean ± SD, n=4*) and analysed by one-way ANOVA followed by Dunnett’s multiple comparison test; **D.** Secreted IL-1β level quantified by ELISA from cells stimulated as in (B) (*mean ± SD, n=6*) and analysed by one-way ANOVA followed by Dunnett’s multiple comparison test; **E.** Immunoblot of NLRP3 (upper panel) and p62/SQSTM1 (lower panel) respectively in macrophages stimulated with LPS/ATP and treated or untreated with different concentrations of tolcapone as indicate din the figure; **F.** Band intensity of NLRP3 normalized to actin for two independent experiments was quantified and represented graphically (*mean ± SD, n=3*); **G.** Band intensity of p62/SQSTM1 normalized to actin for 2 independent experiments was quantified and represented graphically (*mean,* n=2); **H.** Cell stimulated either with DMSO or 10 μM tolcapone as described in (A) were fixed with PFA for 1h at room temperature and stained to label p62/SQSTM1 (green), ASC (red) and IL-1β (gray), **I.** The number of p62 puncta per cell was quantified and represented graphically (*mean± SD, n=56),* **J.** The number of ASC positive cells from an acquired field was also quantified and represented graphically (*mean± SD, n=15),* **K.** Caspase-1 secretion and intracellular level in macrophages stimulated with LPS/ATP and treated or untreated with different concentrations of tolcapone as indicated in the figure; **L.** GSDMD cleavage detected by WB in cells stimulated with LPS and ATP in the presence or absence of different concentrations of tolcapone (FL-full length GSDMD, p30-N-terminus fragment of GSDMD);

Given its consistent inhibitory activity across all tested systems, we next investigated the mechanism by which tolcapone modulates IL-1β export. Tolcapone is a well-characterised inhibitor of catechol-O-methyltransferase (COMT), which catalyses the degradation of catecholamines, including dopamine, epinephrine and norepinephrine, as well as catecholestrogens and other catechol-containing compounds (35).

### Tolcapone impairs caspase-1 activity, limiting GSDMD cleavage and IL-1β secretion

Previous studies have shown that dopamine promotes autophagy-mediated degradation of the NLRP3 (NLR family pyrin domain containing 3) inflammasome via dopamine receptor 1 (DRD1) signaling, thereby limiting NLRP3-dependent inflammatory responses and IL-1β secretion in primary macrophages (36). Based on this, we asked whether tolcapone might indirectly affect inflammasome stability by altering intracellular dopamine levels via COMT. To test this hypothesis, we monitored the expression of NLRP3 and ASC (Apoptosis-associated speck-like protein containing a CARD) in cells treated or not with tolcapone. Tolcapone treatment did not significantly alter the expression of either protein, indicating that its inhibitory effect on IL-1β secretion is unlikely to be mediated through modulation of inflammasome stability via this pathway (Fig. 3E-upper panel, F and S3C-upper panel). Also, we could not observe major differences between the number of ASC positive cells for treated and non-treated cells (Fig. 3H and J). Moreover, the general autophagic flux was not significantly altered by tolcapone treatment as highlighted by the expression level of p62/SQSTM1 (Fig. 3E-lower panel and G) and puncta concentration/cell (Fig. 3H and I) as well as LC3-II/LC3-I ratio (Fig. S3C-lower panel)

Considering that tolcapone did not affect NLRP3 stability and is applied simultaneously with ATP stimulation, we hypothesised it likely acts at later stages of the pathway, prior to cytokine export across the plasma membrane. We therefore assessed the effect of tolcapone on caspase-1 activation and gasdermin D (GSDMD) cleavage. Tolcapone treatment reduced the generation of bioactive caspase-1 (p20) and its release into the extracellular space in a concentration-dependent manner (Fig. 3K). Similarly, cleavage of full length GSDMD to oligomerisation-competent N-terminal fragment (NT) was also inhibited in a dose-dependent manner (Fig. 3L). Together these results suggest that tolcapone impairs caspase-1 activity, thereby limiting GSDMD processing and downstream cytokine maturation and release.

### IL-1β secretion is modulated by tolcapone independently of COMT

Because tolcapone is a COMT inhibitor clinically approved for Parkinson’s disease treatment and considering that it did not affect NLRP3 stability, we next asked whether its effect on IL-1β secretion depends on COMT activity. To address this, we compared tolcapone with two other commercially available COMT inhibitors, opicapone and entacapone, using the same experimental set-up as described in Fig. 3A. Entacapone had no measurable effect on IL-1β secretion for all tested concentrations, as shown by western blotting and ELISA (Fig. 4A–C). Conversely, opicapone had only a mild effect at the highest concentration (10 μM), as determined by Western blotting and ELISA (Fig. 4D–F). Given the differential effects of the two compounds on IL-1β secretion, we next examined whether they influence GSDMD-NT fragment generation in the same samples. Neither entacapone nor opicapone significantly altered GSDMD proteolysis compared to untreated controls (Fig. 4G and H).

**Figure 4.**
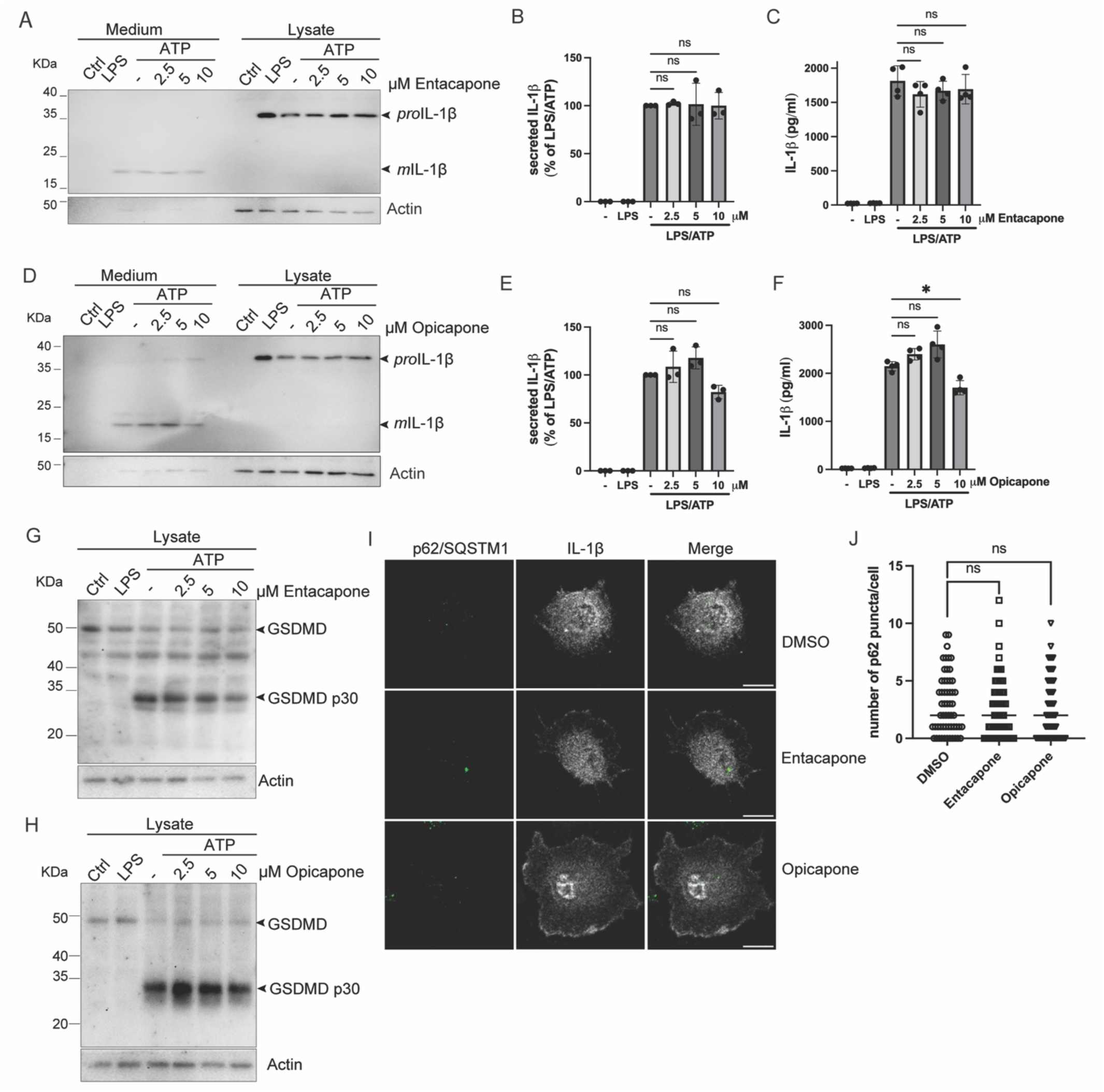
Tolcapone inhibits IL-1β secretion independent COMT. **A.** BMDMs were stimulated as described in Fig. 3A in the presence of a COMT inhibitor, entacapone at the indicated concentrations. Supernatants and lysates were separated by SDS-PAGE, transferred on nitrocellulose membranes and blotted with antibodies against IL-1β or actin; **B.** Band densitometry of three independent experiments presented in (A) (*mean ± SD,* n=3) and analysed by one-way ANOVA followed by Dunnett’s multiple comparison test; **C.** ELISA quantification of secreted IL-1β from cells stimulated as in (A) (*mean ± SD, n=4*) and analysed by one-way ANOVA followed by Dunnett’s multiple comparison test; **D and E.** Western blotting analysis of cells stimulated in the presence of other COMT inhibitor opicapone (D) and band densitometry of three independent experiments (E) (*mean ± SD,* n=3) and analysed by one-way ANOVA followed by Dunnett’s multiple comparison test; **F.** ELISA quantification of secreted IL-1β from cells stimulated as in (D) (*mean ± SD, n=4*) and analysed by one-way ANOVA followed by Dunnett’s multiple comparison test; **G and H.** Immunoblotting analysis of GSDMD proteolysis in macrophages stimulated with LPS/ATP and treated or not with different concentrations of entacapone (G) and opicapone (H) using anti-GSDMD antibodies (FL-full length, p30–N-terminus fragment of GSDMD); **I.** Immunofluorescence analysis of peritoneal macrophages stimulated with LPS for 4h and 20 min with ATP and either DMSO or entacapone or opicapone. Cells were washed, fixed and stained with antibodies for IL-1β (grey), and p62/SQSTM1 (green) as described in Materials and methods; **J.** The number of p62/SQSTM1 puncta per cell was represented graphically (*mean ± SD, n=54*);

We also monitored the number of p62/SQSTM1 puncta/cell as a readout of autophagy activation in cells exposed to the COMT inhibitors and observed no major differences comparted to non-treated samples (Fig. 4I and J). Collectively, these findings suggest that the inhibitory effect of tolcapone on IL-1β secretion is unlikely to depend on COMT activity or to be mediated by differences in autophagy flux.

### LPS-induced IL-1β production is attenuated by tolcapone *in vivo*

Considering the efficient reduction in IL-1β secretion in primary macrophages we next questioned whether tolcapone could attenuate IL-1β-mediated inflammation *in vivo* using a mouse model of LPS-induced endotoxemia (30,37,38). For this mice were administered LPS (1 mg/kg) together with either vehicle (DMSO) or tolcapone (20 mg/kg) and monitored the expression of IL-1β in the liver, spleen and pancreas of treated mice (39–41). We found that administration of LPS resulted in a marked increase in IL-1β expression in the liver as early as 2 h post-injection, which was substantially reduced in tolcapone-treated mice. However, this inhibitory effect was transient, showing a noticeable decline by 4 h and being largely lost by 6 h (Fig. 5A and B). Similar kinetics were observed in the spleen (Fig. 5C and D) and pancreas (Fig. 5E and F). We also assessed whether the inhibitory effect of tolcapone on IL-1β secretion was reflected at the systemic level. Serum IL-1β levels showed a modest reduction in tolcapone-treated mice compared to vehicle controls, however the results displayed a marked variation between experiments (Fig. 5G). Although unexpected, the anti-inflammatory effect of tolcapone is transient with IL-1β levels returning to those observed in untreated animals by 6 h post-treatment, likely reflecting the rapid metabolism of the compound (42,43).

**Figure 5:**
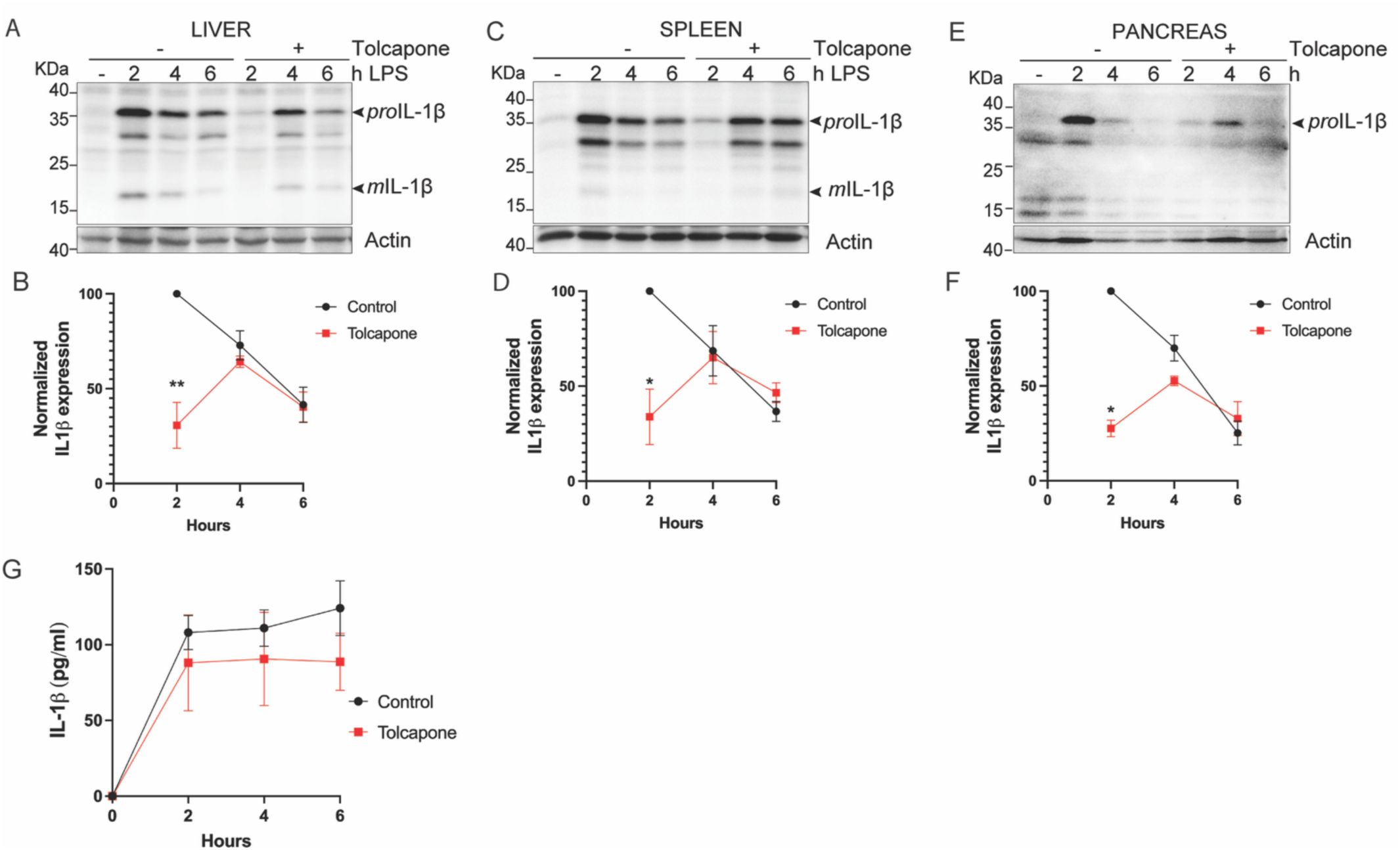
Tolcapone reduces IL-1β after endotoxemia induction. **A.** IL-1β levels, along with actin as loading control, were evaluated by Western blotting for liver extracts from mice stimulated or not with LPS (1mg/kg) to induce sterile sepsis, and tolcapone administration at the indicated time points; **B.** Band intensity quantification of three independent experiments of IL-1β normalized to actin (*mean ± SD, n=3*) and ed by two-way ANOVA followed by Sidak’s multiple comparison test; **C and D.** IL-1β levels in the same conditions as in 4A for spleen (C) and band intensity quantification of three independent experiments of IL-1β bands normalized to actin (*mean ± SD, n=3*) (D) and analysed by two-way ANOVA followed by Sidak’s multiple comparison test; **E and F.** IL-1β levels in the same conditions as in 4A for pancreas (E) and band intensity quantification of three independent experiments of IL-1β bands normalized to actin (*mean ± SD, n=3*) (F) and analysed by two-way ANOVA followed by Sidak’s multiple comparison test; **G.** Serum level of IL-1β from mice treated as in (G) quantified by ELISA (*mean ± SD,* n=3);

### Tolcapone inhibits the secretion of inflammasome-dependent cytokines

Next we questioned whether tolcapone modulates the secretion of inflammatory cytokines beyond IL-1β and analysed two additional IL-1 family members: IL-18, an inflammasome-dependent cytokine that shares the same export pathway, and IL-1α, an atypical cytokine found in both soluble and membrane-bound forms and secreted via a distinct pathway (3,44–46). When analysing the effect in stimulated primary macrophages, we observed tolcapone induced a concentration-dependent decrease in secreted IL-18 (Fig. 6A-upper panel and quantification in B). In contrast, IL-1α expression remained largely unaffected intracellularly, with no detectable secretion under the same experimental conditions (Fig. 6A middle panel). These findings indicate that tolcapone preferentially inhibits the secretion of inflammasome-dependent cytokines, such as IL-1β and IL-18, while having minimal impact on cytokines that follow distinct export pathways.

**Figure 6:**
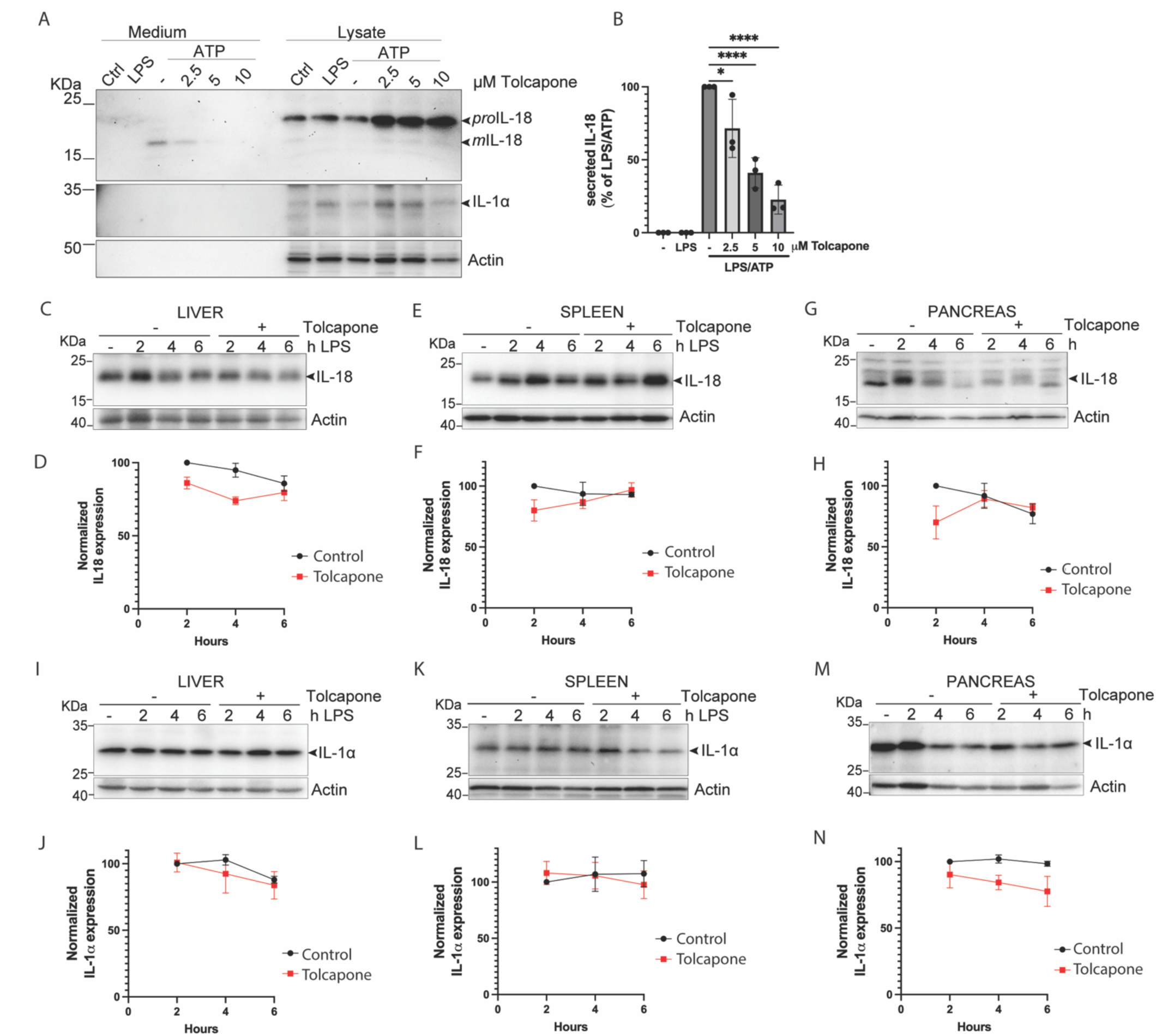
The secretion of IL-18, but not IL-1α is modulated by tolcapone. **A.** BMDMs were stimulated as described in Fig. 3A in the presence of tolcapone at the indicated concentrations. Supernatants and lysates were separated by SDS-PAGE, transferred on nitrocellulose membranes and blotted with antibodies against IL-18, IL-1α or actin; **B.** Band intensity quantification of three independent experiments IL-18 expression normalized to actin (*mean ± SD,* n=3) and analysed by one-way ANOVA followed by Dunnett’s multiple comparison test; **C.** IL-18 levels, along with actin as loading control, were evaluated for liver extracts from mice stimulated or not with LPS (1 mg/kg) to induce endo, and tolcapone (20 mg/kg) administration at the indicated time points; **D.** Band intensity quantification of three independent experiments for IL-18 expression normalized to actin (*mean ± SD, n=3*) and analysed by two-way ANOVA followed by Sidak’s multiple comparison test; **E and G.** IL-18 levels in the same conditions as in (C) for spleen (E) and pancreas (G); **F and H.** Band intensity quantification of three independent experiments normalized to actin for spleen (F) and pancreas (H) (*mean ± SD, n=3*) and analysed by two-way ANOVA followed by Sidak’s multiple comparison test; **I.** IL-1α levels, along with actin as loading control, were evaluated for liver extracts from mice stimulated or not with LPS (1mg/kg) to induce endotoxemia and tolcapone administration at the indicated time points; **J.** Band intensity quantification of three independent experiments of IL-1α bands normalized to actin (*mean ± SD, n=3*) and analysed by two-way ANOVA followed by Sidak’s multiple comparison test; **K and L.** IL-1α levels in the same conditions as in (I) for spleen (K) and band intensity quantification of three independent experiments of IL-1α bands normalized to actin (L) (*mean ± SD, n=3*) and analysed by two-way ANOVA followed by Sidak’s multiple comparison test; **M and N.** IL-1α levels in the same conditions as in (I) for pancreas (M) and band intensity quantification of three independent experiments of IL-1α bands normalized to actin (N) (*mean ± SD, n=3*) and analysed by two-way ANOVA followed by Sidak’s multiple comparison test;

Considering our results showed that tolcapone reduced IL-18 secretion *in vitro*, we next questioned whether similar effects could be observed *in vivo*. We therefore analysed IL-18 expression in the liver, spleen and pancreas of mice treated as described above in Fig. 5. Tolcapone treatment resulted in a modest reduction in IL-18 levels in the liver at 2 h post-injection, which was maintained until 4 h, albeit less prominently than for IL-1β (Fig. 6C and D). In the spleen (Fig. 6E and F) and pancreas (Fig. 6G and H), a transient reduction in IL-18 expression was observed at 2 h, with no significant differences detected at later time points. Notably, by contrast to IL-1β, IL-18 could be detected in both non-stimulated cells as well as in the organs of non-treated mice, and its expression was fairly stable within the 6h timeframe compared with IL-1β which shows a more dynamic expression, potentially explaining the different signal amplitude as response to tolcapone treatment (47–49).

Next, we assessed the effect of tolcapone on IL-1α expression *in vivo*. As expected, tolcapone treatment did not significantly alter IL-1α levels in the liver compared to vehicle controls with no notable changes observed across time points (Fig. 6I and J). Similarly, no major differences were observed for IL-1α expression in the spleen (Fig. 6K and L). In the pancreas, a modest reduction in IL-1α expression was detected following tolcapone administration, although the extensive variability between experiments restricted our ability to draw definitive conclusions (Fig. 6 M and N). Overall, these findings suggest that tolcapone preferentially targets inflammasome-dependent cytokines, attenuating IL-18 production *in vivo,* additional to IL-1β, while having minimal impact on IL-1α regulation.

## Discussion

Inflammasome-driven cytokine release and lytic cell death are emerging as key effectors of inflammatory disorders across diverse pathologies (50–53). Within the IL-1 family, IL-1β acts as a ubiquitous mediator of inflammatory responses that is unconventionally secreted by activated macrophages (5). Although attractive for therapeutic intervention, IL-1β release depends on a coordinated cascade: inflammasome activation, caspase-1 catalytic activity, and subsequent GSDMD pore formation. Consequently, identifying compounds that modulate endogenous IL-1β secretion in physiologically relevant context is challenging. Moreover, the limited repertoire of approved therapies further support the search for new molecules targeting this pathway to control the IL-1β-induced inflammatory phenotype (7,53).

To address this gap, we generated a scalable, screening-compatible reporter cell line that quantitatively monitors endogenous IL-1β secretion in response to canonical inflammatory stimuli such as LPS and ATP. This feature is particularly important, as it facilitates monitoring of IL-1β release in a physiologically relevant context and it allows us to clearly discriminate between transcriptional and secretion regulators. Current therapies often rely on blocking extracellular IL-1β or its receptor, assuming that disease propagation is exclusively driven by autocrine or paracrine signaling. By instead targeting the intracellular events regulating secretion, we can inherently attenuate downstream IL-1 receptor signaling while preventing the concomitant release of other pro-inflammatory mediators. Thus, our system provides a highly robust platform for the discovery of mechanistically distinct anti-inflammatory molecules.

Exploiting this optimized screening platform, we identified and validated several compounds that efficiently blocked IL-1β secretion, including molecules with previously established anti-inflammatory properties (29–32). Among these, the COMT inhibitor tolcapone emerged as consistently inhibiting IL-1β secretion throughout the screening pipeline, which we subsequently validated in primary macrophages and an *in vivo* model of endotoxemia. Tolcapone is clinically utilized as an adjunct to levodopa for Parkinson’s disease (with and IC50 of 3nM in the brain and 795 nM in the liver) and although it its clinical use has been limited by hepatotoxicity its efficacy was superior compared to the newer-generation inhibitors and still prevails in therapy (55–58). Our results suggest that, in macrophages, tolcapone supresses caspase-1 activity, thereby impairing downstream events like GSDMD cleavage, plasma membrane pore formation and the subsequent processing and release of IL-1β. Surprisingly, newer-generation COMT inhibitors did not phenocopy this caspase-1 inhibition or the downstream effects. Moreover, this suppression was independent of inflammasome stability linked to inactive COMT, differentiating our findings from prior reports suggesting dopamine-dependent NLRP3 degradation (36). Together, these data strongly indicate that tolcapone inhibits caspase-1 through a previously unrecognized, COMT-independent mechanism.

Given the pivotal role of GSDMD-mediated pyroptosis in driving tissue damage across diverse pathologies, targeting gasdermin cleavage and pore formation holds immense therapeutic promise (50,59,60). Our study demonstrates that tolcapone significantly impaired GSDMD cleavage in primary macrophages, by contrast with the newer-generation COMT inhibitors, and represents a potential candidate for broader anti-inflammatory interventions targeting both cytokine release and inflammatory cell death. Our results also suggest this effect is selective, tolcapone potently suppressed the release of both IL-1β and IL-18, however it exerted minimal influence on IL-1α. This distinct profile aligns with the paradigm that these IL-1 family members utilize related, yet mechanistically divergent, secretion routes, further confirming that tolcapone specifically affects the inflammasome/caspase-1/GSDMD pathway (14).

Although direct inhibition of inflammatory caspases represents an attractive therapeutic strategy, the clinical translation of these candidates has been hindered by dose-limiting toxicities, including the positive control used in our assay, AC-YVAD-MCK (61). Because tolcapone is already FDA-approved, its newly discovered off-target anti-inflammatory activity presents an attractive opportunity for drug repurposing, potentially bypassing limitations associated with novel caspase-1 inhibitors. Our *in vivo* data confirms that tolcapone can limit acute septic response after LPS-induced endotoxemia. Nevertheless, its effect appears transient, likely reflecting the relatively short half-life of the compound. Interestingly, this short *in vivo* half-life could be strategically advantageous for acute inflammatory settings, allowing for a transient suppression of the cytokine storm without inducing a prolonged blockade of caspase-1, which remains essential for baseline innate immune defence.

While the LPS endotoxemia model effectively recapitulates the acute cytokine storm of early-stage sepsis, future investigations utilizing polymicrobial infection models, such as caecal ligation and puncture (CLP), are required to fully outline the efficacy of tolcapone for the multiphasic pathophysiology of clinical sepsis. Furthermore, identifying the direct molecular target of tolcapone, whether it interacts directly with caspase-1 or modulates an immediate upstream regulator, remains an essential next step. Ultimately, our findings establish a powerful, scalable platform for discovering modulators of unconventional secretion and highlight tolcapone as a promising, COMT-independent inhibitor of inflammasome-driven acute septic response.

## Materials and methods

### Chemicals and reagents

The FDA-approved Drug Library (L4200) containing 1398 compounds, used for screening with the IL-1β HiBiT reporter cell line was from TargetMol. Individual compounds used for validation experiments: tolcapone (HY-17406), teriflunomide (HY-15405), nimodipine (HY-B0265), Ac-YVAD-cmk-Caspase-1 inhibitor II (HY-16990), were from MedChemExpress. Blotto, non-fat dry milk (sc-2324), Tris (sc-3715), Tween 20 (sc-29113), glycine (sc-29096) were from Santa Cruz Biotechnology; all other chemicals were from Sigma Aldrich, unless otherwise specified.

### Mice

C57BL/6J mice were maintained under specific pathogen-free conditions at 22±2°C and given access to food and water ad libitum. Animals of both sexes were used for all experiments and mice with age between 10-16 weeks were used for experiments, no criteria for inclusion or exclusion have been applied. Experiments were performed in the Animal Facility of the “Victor Babeș” National Institute of Pathology (Bucharest, Romania) and were approved by the Ethics Committee of Victor Babeș National Institute of Pathology (protocol code 66/14.09.2019) and by the National Sanitary Veterinary and Food Safety Authority, Bucharest, Romania (project authorization no. 476/04.11.2019).

### Cell culture

J774A.1 macrophage cell line (ATCC-TIB-67) was purchased from ATCC. Cells were grown in DMEM with Glutamax (Gibco-31966-047)) supplemented with 10% FBS (Gibco-10437-28), 1% Penicillin/Streptomycin (Gibco-1540-122) and were maintained at 37C, with 5% CO2 and 95% relative humidity. Cells were cultivated according to the ATCC instructions and always maintained at sub-confluent density. Primary macrophages were grown in DMEM/F12 with Glutamax (Gibco-21331-028), supplemented with, 10% serum, 10% L929 supernatant, 1% Penicillin/Streptomycin. HeLa cells stably expressing HiBiT-tagged cargoes were grown in DMEM with Glutamax (Gibco-31966-047) with 10% FBS (Gibco-10437-28), 1% Penicillin/Streptomycin (Gibco-1540-122).

Peritoneal macrophages (PMs) and bone marrow-derived macrophages (BMDMs) were isolated and cultivated as previously described (23). **PMs:** cells were collected by lavage of mouse peritoneal cavity with 5 ml of PBS-4% FBS, centrifuged at 1000rpm for 10 min at room temperature, resuspended in complete medium and counted. 2×10^5^ cells were seeded onto coverslips, allowed to adhere for 4-6h and afterwards the medium was replaced. Three days later, the cells were processed for immunofluorescence experiments as described below. **BMDMs:** Bone cavity from tibias and femurs were flushed with complete medium to collect the bone marrow. The tissue was dislodged, passed through a cell strainer (70 mm mesh) and centrifuged to remove cell debris. 8-10×10^6^ cells were seeded onto 100mm dishes in 15 mL of medium and grown at 37C. Four days later, equal amount of medium was added to the dish and maintained in culture for additional days. Cells were scraped, counted and seeded in 12 well plates and used for stimulation 2-3 days later.

### IL-1β HiBiT CRISPR/Cas9 genome editing and cell line generation

The genome editing strategy was designed to fuse a flexible spacer (GGsGG) and HiBiT tag (Promega) to the C-terminus of the IL-1β-encoding gene. Ribonucleoprotein (RNP) complexes were prepared as previously described using the Alt-R CRISPR-Cas9 System components comprising Alt-R custom crRNA, tracrRNA fluorescently labelled with ATTO 550 (IDT-1075927), recombinant Cas9 (IDT-1081060), and the Ultramer single-stranded oligodeoxynucleotide donor DNA (ssODN) (26). The crRNAs were designed using Chop-Chop program (http://chopchop.cbu.uib.no/) and all sequences are listed in Table 1. Briefly, equimolar amounts of crRNA and tracrRNA were mixed in Duplex buffer (IDT) to a final concentration of 12 μM and annealed to form the guide RNA (gRNA) duplex by heating at 95°C for 5 min and cooling at room temperature. Then, 50 pmol of Cas9 enzyme were slowly mixed with 60 pmol gRNA duplex to form RNP complexes for 20 min at room temperature. The RNP complexes and the ssODN donor DNA sequences were electroporated to J774A.1 cells using the Neon NxT electroporator (Thermo Scientific) with the Neon Transfection System 10 μL kit (MPK1025) following the manufacturer’s instructions. Briefly, 250 000 cells were resuspended in Buffer R, combined with the ssODN-RNP complex prepared in Opti-MEM media (Gibco-31985-070) and electroporated with the following settings: voltage 1700, width 20ms, pulses 2. Next, the cells were plated in complete growth media and let to recover for 48h. Single cell sorting based on the tracrRNA fluorescence (ATTO 550) and individual clones were tested for HiBiT luminescence and PCR genotyping.

PCR genotyping was done with the Mouse direct PCR kit (Bimake B40015), using 200,000 cells per 25 μL lysis buffer mix according to the manufacturer’s instructions. The presence of a 284bp amplicon corresponding to insertion of HiBiT sequence could be observed for positive clones. Homozygous clones were selected for further validation by luminescence assay. The positive clones were sequenced to confirm the presence of HiBiT.

**IL-1β Secretion Assay:** J774A.1 and J774A.1-IL-1β HiBiT cells were stimulated with 0.5 μg/ml LPS (ALX-581-007-L002, EnzoLifeSciences) for about 16h in complete medium; next day cells were washed with PBS and stimulated for 50 min with 2 mM ATP (A9187, Sigma), in Opti-MEM. Supernatants were collected, spun down to remove cell debris and processed immediately for luminescence assay or were stored at –80C for ELISA or Western blotting analysis.

BMDMs were differentiated for 7 days in the presence of growth factors (L929 conditioned medium), detached and seeded 2*10^5^/12w plates. 3 days later cells were stimulated with LPS 0.5 μg/mL in complete medium for 4h, washed with PBS and incubated with 2 mM ATP/0.5 μg/mL LPS in physiological saline solution (PSS-147 mM NaCl, 10 mM HEPES pH 7.4, 13 mM glucose, 2 mM CaCl2, 1 mM MgCl2, 2 mM KCl) (62). Supernatants were collected, centrifuged and aliquoted for either ELISA or WB, and cells were lysed in buffer containing 1% Triton-X100 (20 mM Tris-HCl pH 7.4, 150 mM NaCl, 1 mM CaCl2, 1 mM MgCl2, 1% Triton-X100) supplemented with protease inhibitor cocktail (04693116001 Roche) 1x in final volume, 10 mM NaF, 1 mM Sodium orthovanadate, 1 mM b-glycerophosphate, 1 mM EDTA and 20 mM NEM (N-ethylmaleimide) for 30 min on ice. The lysates were cleared out by centrifugation at 10000xg, for 20 min.

### Large-scale screening and luminescence measurement

The amount of IL-1β-HiBiT secreted, under stimulated conditions in the extracellular medium was measured using Nano-Glo HiBiT Extracellular Detection System kit (N2421-Promega). This system contains the HiBiT complementary Large BiT (LgBiT) subunit of the NanoBiT Luciferase and the furimazine substrate, necessary to detect the enzyme activity; the luminescencent signal is proportional with the amount of IL-1β-HiBiT secreted in the medium. J774.A1-IL-1β-HiBiT cells were seeded in 96 well plates, let to adhere for approximately 24h then stimulated with 0.5 ug/mL LPS for 16 h in complete growth medium. Next day the medium was replaced with OptiMEM containing 0.5 μg/ml LPS and 2mM ATP supplemented with either vehicle control (DMSO) or inhibitors at 20 μM concentration, for 50 min. Freshly collected cell supernatants were mixed in a 1:1 ratio (v/v) with the Nano-Glo HiBiT Extracellular Reagent. The reaction mixture for each sample was prepared in opaque white 96-well tissue culture plates, and the resulting Relative Luminescence Units (RLU) were measured using a spectrophotometer (Berthold Technologies Mithras-LB940) with an integration time of 1 second. The RLU values were corrected by subtracting the background values of the control under non-stimulated conditions and graphically represented.

Primary screening: Supernatants of stimulated cells and treated with DMSO, Ac-YVAD-CMK or 20 µM individual compounds of the FDA library (3 replicates per sample), were mixed at 1:1 ratio (v/v) with the Nano-Glo HiBiT Extracellular Detection mix and afterwards luminescence was measured as described above.

Secondary screening: Supernatants of stimulated cells (4 replicates for each compound) of DMSO-treated controls and the top 74 compounds (selected based on the RLU log2 ratio of drug/DMSO≤-2,320) at 10 µM concentration were used for detection of secreted IL-1β-HiBiT by luminescence measurement as described above.

<u>Dose response calculation:</u> J774A.1-IL-1β-HiBiT cells were stimulated with 0.5 ug/ml LPS for 16h in complete medium; afterwards the cells were washed with PBS and stimulated for 50 min with 0.5 ug/ml LPS and 2 mM ATP in Opti-MEM in the presence of different concentrations of the indicated chemicals (three-fold dilution from 900 µM to 30 nM). Supernatants were collected and processed immediately for luminescence assay. The obtained RLU normalized to background values obtained in non-stimulated conditions (LPS treated) were graphically represented in Graph Pad Prism 11 as logarithmic dose response curve, using a non-linear regression function to fit the data.

### HiBiT Secretion Assay in HeLa cells

HeLa cells expressing the HiBiT tagged reporter were seeded in a 96-well plates 96F TC (Black, Clear bottom, PerkinElmer #6005182) and let to adhere for approximately 24h. Cells were then washed three time with PBS and incubated in fresh complete medium for the indicated time. As previously described in HeLa cells (33,34), the split luciferase complementation assay was performed from 100 μL of cell culture medium using the Nano-Glo HiBiT Extracellular Detection System Kit (#N2421, Promega), and from 100 μL of cells permeabilized using the Nano-Glo HiBiT Lytic Detection System Kit (#N3040, Promega) according to manufacturer’s instructions. Luminescent signal was measured using a Tecan Microplate reader Infinite 500 with an integration time of 0.5-2 seconds. The obtained RLU values for each condition were normalized to values obtained in control condition (DMSO at t=30min) and afterwards graphically represented.

Lactate dehydrogenase (LDH) release was monitored from cell culture media collected at the indicated times to assess the extent of plasma membrane damage and cell death using a commercially available LDH assay kit (Abcam, #ab102526), according to the manufacturer’s recommendations.

### ELISA

IL-1β secreted levels were determined using Mouse DuoSet ELISA kit (DY401-05), following the manufacturer’s instructions. Medium fractions were thawed on ice and 4-10% of the collected media was used for sandwich ELISA in 2 replicates for each sample. The final absorbance values were read at spectrophotometer (FLUOstar Omega, BMG Labtech). The normalized absorbance values at 450 nm were used to estimate the concentration of secreted cytokine.

### Western blotting

For SDS-PAGE and Western blotting analysis of secreted and intracellular expression of IL-1β (tagged and non-tagged version) 1/5 (v/v) of cell lysates were separated along corresponding volumes of supernatants under denaturing conditions. For total lysates analysis from cells and tissues extracts, equal amounts of protein were separated on 8-12% acrylamide gels, depending on the protein of interest and then transferred onto nitrocellulose or PVDF membranes (cat no: 10600001 and cat no: 10600023, Amersham).

*Antibodies-based detection:* the membranes were blocked in 5% BSA in TBS-Tween 0.1% (TBS 50 mM Tris-Cl, pH 7.5, 150 mM NaCl) and probed with the indicated primary antibodies overnight at 4°C in 5% BSA in TBS-Tween 0.1%. Next day, HRP coupled secondary antibodies (Jackson ImmunoResearch) were added for 1h at room temperature in 5% BSA in TBS containing 0.1% Tween and visualization was made using Immobilon Crescendo Western HRP substrate (cat no: WBLUR0500, Merck-Millipore).

*Detection with Nano-Glo HiBiT Blotting antibody free System kit (Promega –N2410):* nitrocellulose membranes were incubated in 0.1% TBS-Tween for 30 minutes to 1hour to expose HiBiT, allowing LgBiT binding after an overnight incubation at a 1:200 dilution in 1× Nano-Glo Blotting Buffer. The resulting LgBiT/HiBiT complex generates a luminescent signal upon addition of the Nano-GloLuciferase Assay containing the furimazine substrate, which was detected using a ChemiDoc Imaging system (Bio-Rad).

### Immunofluorescence microscopy

J774A.1-IL-1β-HiBiT cells were grown onto glass coverslips and stimulated with 0.5 μg/mL LPS for 16h. Next day the cells were stimulated for 30 min with LPS (0.5 μg/mL) and 2mM ATP, fixed by incubation with 1% PFA (paraformaldehyde) in PBS (Phosphate buffered saline-13mM NaCl, 2.7 mM KCl, 10 mM Na2HPO4, 1.8 mM KH2PO4) for 1h. Peritoneal macrophages were cultivated onto cover glasses for 3 days, stimulated with LPS (0.5 μg/ml) for 4h and ATP (2mM) for 15 min, washed with PBS and then fixed by incubation with 1% PFA in PBS for 1h.

Subsequently, the samples were permeabilized for 3 min with 0.0125% digitonin in blocking buffer (2% horse serum in PBS), incubated with blocking buffer for 2h and then with primary antibodies overnight at the indicated dilutions in humidified atmosphere. Next day, the coverslips were washed and incubated with the corresponding secondary antibodies or combination of secondary antibodies and fluorescent dyes for 30 min at room temperature. Samples were washed extensively and subsequently mounted on glass slides. Image acquisition was made using Zeiss LSM confocal microscope (63x objective, PA, 1.4 NA) using Zen software.

### LPS-induced endotoxemia in mice and sample collection

Mice were weighted the day of the experiment and similar weights animals were paired for each time point. Tolcapone (20 mg/kg) dissolved in a mix of DMSO: PEG300/PEG400:Tween-80:saline (10% DMSO:40% PEG300/PEG400:5% Tween-80:45% saline) or vehicle control were injected intraperitoneally (IP) to mice and within 5 minutes the LPS (1mg/kg) was also injected IP to all mice. Blood was collected by cardiac puncture under anaesthesia (ketamine/acepromazine cocktail; ketamine 100 mg/kg, ketamin 10%, Medistar Arzneimittelvertrieb Gmbh, Ascheberg, Germany; acepromazine 5 mg/kg, Calmivet Solution Injectable Acepromazine 5 mg, Vétoquinol SA, Lure, France) and organs (liver, spleen and pancreas) as well as peritoneal macrophages were collected postmortem for mice exposed to LPS and drugs/DMSO for 2, 4 and 6h. Serum was isolated from the blood fraction, aliquoted and stored at –80°C for further processing.

*Total protein extraction from organs:* fraction of the organs were minced in RIPA lysis buffer (25mM Tris HCl, pH 7.6, 150mM NaCl, 1% NP-40, 1% Sodium Deoxicholate, 0.1%SDS) supplemented with protease inhibitor cocktail 1x in final volume and phosphatases inhibitors:10 mM NaF, 1 mM Sodium orthovanadate, 1 mM b-glycerophosphate, at 20 ug/ml and extracted using an ultrasonic homogenizer (Bandelin Sonoplus). The extraction was performed on ice, with following settings: 6 pulses x3 seconds, 30% power for liver and 10 pulses x 3 seconds, 65% power for spleen and pancreas. Extracts were cleared out by centrifugation at 20000xg for 20 min at 4C. Equal amounts from each sample were separated by SDS PAGE, in reducing conditions, transferred onto nitrocellulose or PVDF membranes and probed with the indicated antibodies.

### Statistical analysis

*High-throughput Screening:* <u>Primary screening:</u> For each compound from the individual 96-well plates, mean of 3 technical replicates, the corresponding coefficient of variation were calculated and the ratio Drug /DMSO. The obtained values are plotted as log Drug/DMSO vs CV. Compounds exhibiting a ratio drug DMSO, less than or equal to 0.2 (log2:-2,32) and a CV below 30% in this primary screen were preselected for secondary screening.

Secondary screening, mean of 4 biological replicates and SD were calculated for each compound and a threshold of 0.75 (log2: –0.4775) Drug/DMSO ratio was established.

*Statistical analysis for ELISA or band densitometry:* All values of ELISA, luminescence and WB quantification are expressed as the mean ± SD unless otherwise specified in the figure legend. For the ELISA experiments and WB band densitometry statistical analysis was performed using one-way or two-way ANOVA (GraphPad Software Prism 11) for multiple groups with the recommended correction for each method. P values < 0.05 (*) were considered significant.

## Acknowledgements

We thank dr. Ombretta Foresti (Centre for Genomic Regulation, Barcelona, Spain) for critically reading this manuscript and the members of our laboratory for helpful discussions. We thank dr. Mihaela Uta (Haematology Research Laboratory, Fundeni Clinical Institute), dr. Livia Sima, and drd. Cosmin Trif (Institute of Biochemistry of the Romanian Academy, Bucharest, Romania) for valuable discussion and technical assistance.

## Author Contribution Statement

GC: conceptualization, methodology, investigation, visualization, writing – original draft, writing – review & editing,

SG, GI, JV: conceptualization, methodology, investigation, writing – review & editing,

CVAM, CP visualization, data curation, writing – review & editing

SMP, resources, methodology, writing – review & editing

AB, visualization, data curation, writing – review & editing,

MCB: conceptualization, methodology, investigation, visualization, funding acquisition, supervision, writing – original draft, writing – review & editing,

## Ethics Statement

All procedures were performed in compliance with the relevant guidelines and regulations, as approved by the Ethics Committee of Victor Babeș National Institute of Pathology (protocol code 66/14.09.2019) and by the National Sanitary Veterinary and Food Safety Authority, Bucharest, Romania (project authorization no. 476/04.11.2019).

## Data Availability Statement

The authors confirm that the data supporting the findings of this study are available within the article or its supplementary materials. Correspondence and material requests should be addressed to Marioara Chiritoiu-Butnaru.

## Figure legends

**Supplementary figure 1:**
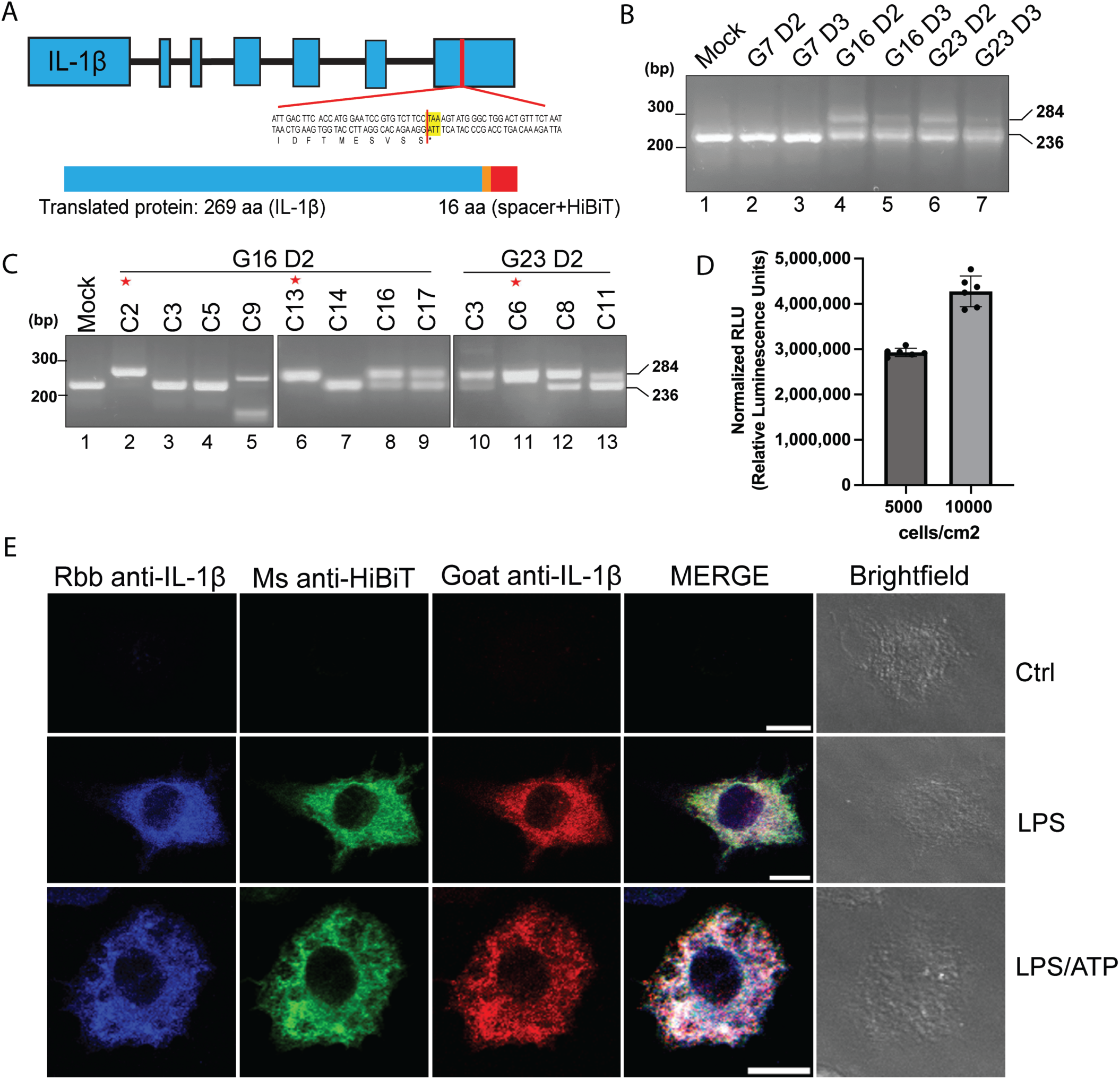
A. Schematic diagram of IL-1β-HiBiT translated sequence showing the spacer and HiBiT insertion at the C-terminus of the IL-1β translated gene; B. PCR genotyping of electroporated cells. 48h after electroporation, the cells were harvested and lysed for genomic DNA extraction. The sequence encoding for exon 7 was amplified by PCR with specific primers (see oligonucleotides table) and visualized by agarose gel electrophoresis. Lower band (236 bp) indicates endogenous fragment, upper band (284 bp) indicates HiBiT inserted fragment; C. Single cell clones were expanded, and an aliquot of cells were used for PCR genotyping and visualized by agarose gel electrophoresis; D. Cells were seeded at different densities 5000 and 10,000 cells/cm^2^, stimulated with LPS and ATP and their extracellular luminescence was evaluated. Values of three independent experiments each with two biological replicates/experiment are represented graphically (*mean ± SD,* n=6); E. Immunofluorescence microscopy for J774.A1 expressing IL-1β-HiBiT clone with three antibodies recognizing HiBiT or IL-1β (rabbit anti-IL-1β-blue, mouse anti-HiBiT-green, goat anti-IL-1β-red);

**Supplementary figure 2:**
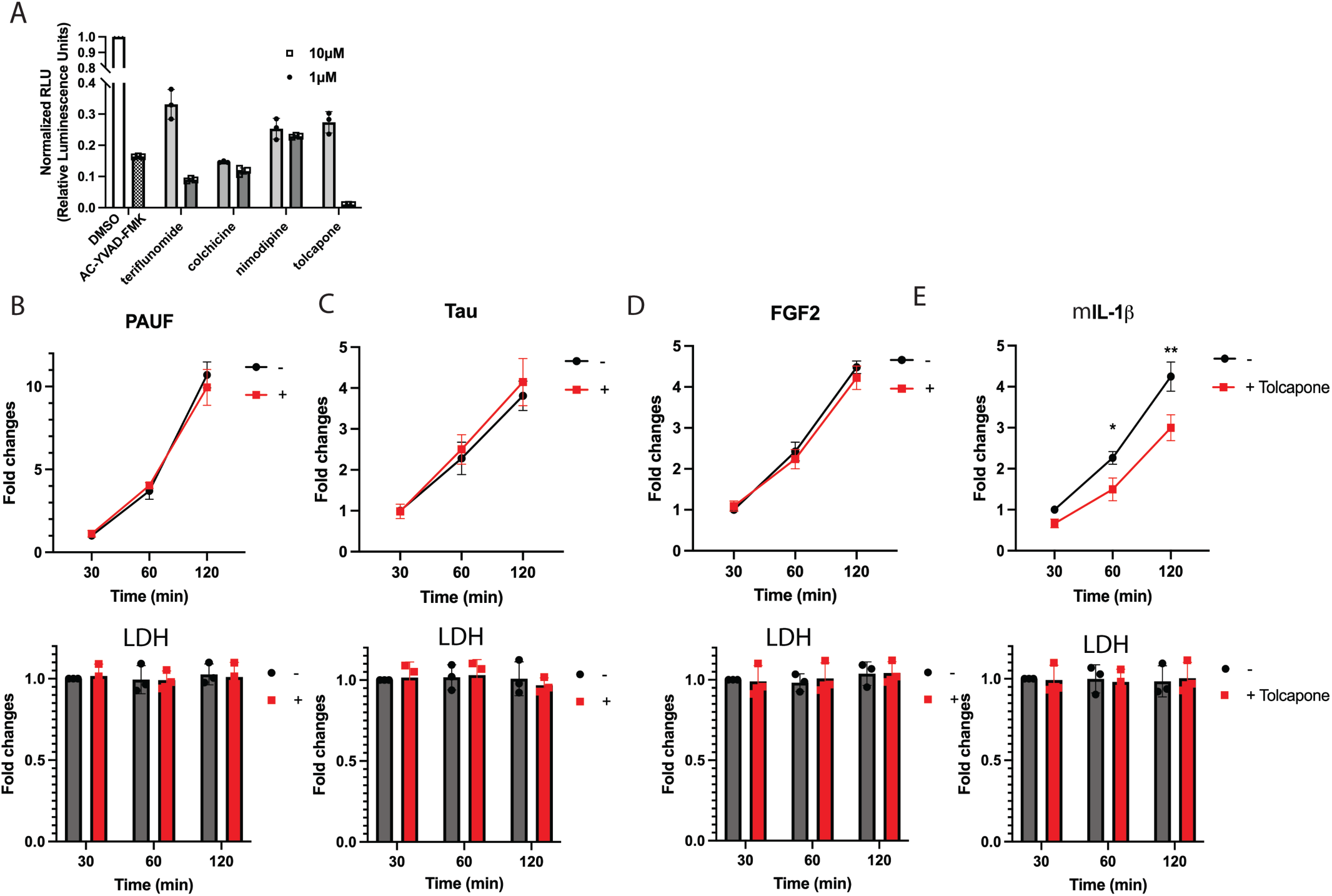
A. Cells were seed in 96 well plates and stimulated as described for screening purposes (figure 1A) in the presence or absence of the caspase-1 inhibitor Ac-YVAD-CMK (10 μM) as positive control and the four selected drugs: teriflunomide, colchicine, nimodipine and tolcapone at two concentrations 10 μM and 1 μM; B. B to E. Split luciferase complementation assay in HeLa cells expressing either PAUF-HiBiT (B), Tau-HiBiT (C), FGF2-HiBiT (D), and IL-1β-HiBiT (E) incubated in complete medium in a 120-min time course. The ratio of medium to lysate luminescence was quantified and the values for each condition were normalized to t=30 min (*mean of n = 3 ± SD; **p<0.01* vs. DMSO condition). LDH release was also monitored from each medium fraction (*mean of n = 3 ± SD*) and analysed by two-way ANOVA followed by Sidak’s multiple comparison test. The specificity of the assay was validated by the absence of a difference in LDH release between all conditions;

**Supplementary figure 3:**
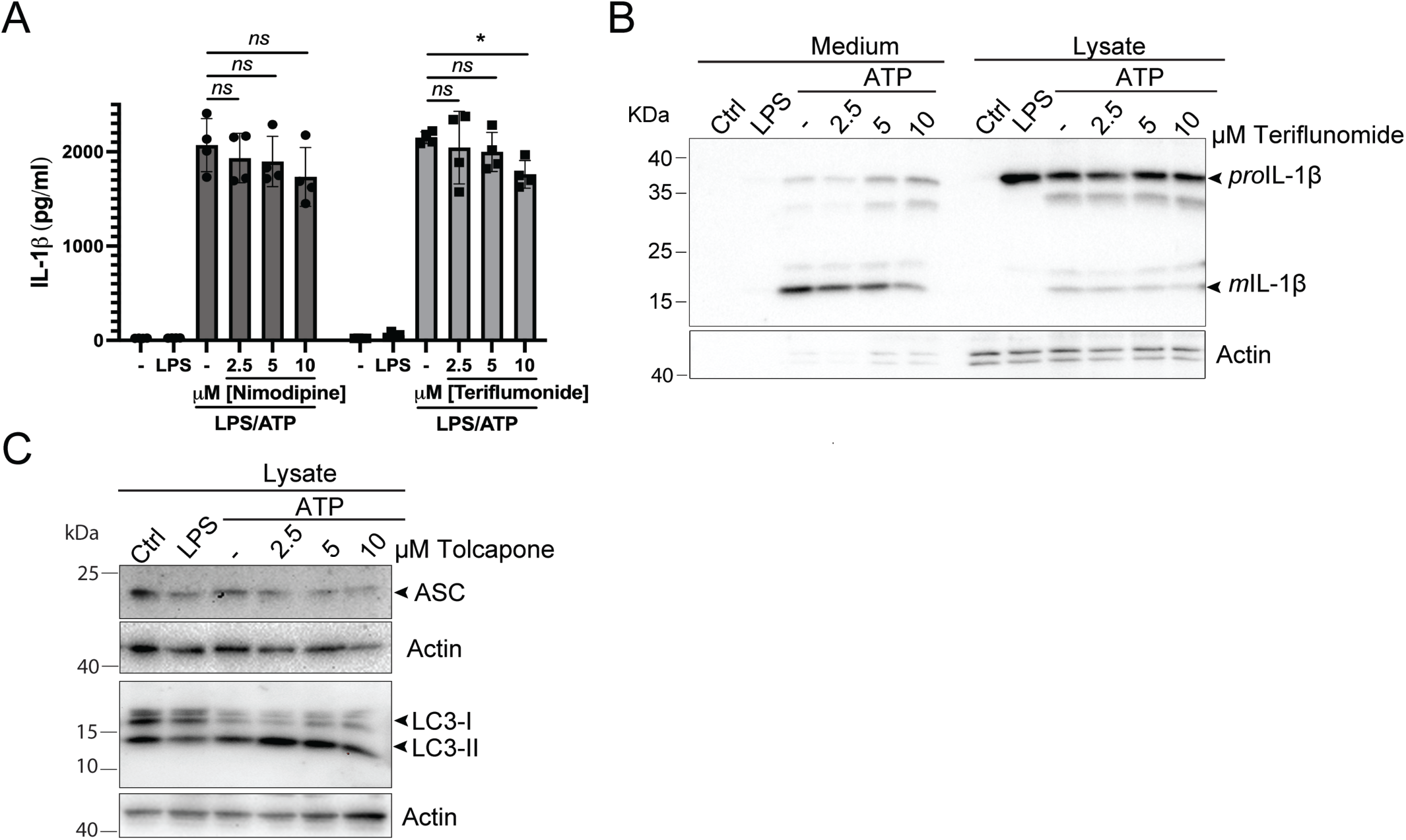
A. Secreted IL-1β level quantified by ELISA from BMDMs stimulated in the presence of different concentrations of nimodipine and teriflunomide (*mean ± SD, n=*4, \**p<0.05*) and analysed by one-way ANOVA followed by Dunnett’s multiple comparison test; B. Western blotting evaluation of IL-1β levels, along with actin as loading control, for BMDMs stimulated in the presence of different concentrations of teriflunomide; C. Western blotting analysis of ASC (upper panel), LC3I/LC3II (lower panel) and actin as loading control for BMDM treated with different tolcapone concentrations;

## Reference

1. Placha D, Jampilek J. Chronic Inflammatory Diseases, Anti-Inflammatory Agents and Their Delivery Nanosystems. Pharmaceutics. 2021 Jan 6;13(1):64. doi:10.3390/pharmaceutics13010064

2. Chen L, Deng H, Cui H, Fang J, Zuo Z, Deng J, et al. Inflammatory responses and inflammation-associated diseases in organs. Oncotarget. 2018 Jan 23;9(6):7204–18. doi:10.18632/oncotarget.23208

3. Gabay C, Lamacchia C, Palmer G. IL-1 pathways in inflammation and human diseases. Nat Rev Rheumatol. 2010 Apr;6(4):232–41. doi:10.1038/nrrheum.2010.4

4. Auron PE, Webb AC, Rosenwasser LJ, Mucci SF, Rich A, Wolff SM, et al. Nucleotide sequence of human monocyte interleukin 1 precursor cDNA. Proc Natl Acad Sci U S A. 1984 Dec;81(24):7907–11. doi:10.1073/pnas.81.24.7907 PubMed PMID: 6083565; PubMed Central PMCID: PMC392262.

5. Dinarello CA. Interleukin-1 in the pathogenesis and treatment of inflammatory diseases. Blood. 2011 Apr 7;117(14):3720–32. doi:10.1182/blood-2010-07-273417

6. Evavold CL, Kagan JC. Diverse Control Mechanisms of the Interleukin-1 Cytokine Family. Front Cell Dev Biol. 2022 Jun 27;10:910983. doi:10.3389/fcell.2022.910983

7. Hommel U, Hurth K, Rondeau JM, Vulpetti A, Ostermeier D, Boettcher A, et al. Discovery of a selective and biologically active low-molecular weight antagonist of human interleukin-1β. Nat Commun. 2023 Sep 7;14(1):5497. doi:10.1038/s41467-023-41190-0

8. Dinarello CA, Simon A, Van Der Meer JWM. Treating inflammation by blocking interleukin-1 in a broad spectrum of diseases. Nat Rev Drug Discov. 2012 Aug;11(8):633–52. doi:10.1038/nrd3800

9. Mai W, Liao Y. Targeting IL-1β in the Treatment of Atherosclerosis. Front Immunol. 2020 Dec 10;11:589654. doi:10.3389/fimmu.2020.589654

10. Eder C. Mechanisms of interleukin-1β release. Immunobiology. 2009 Jul;214(7):543–53. doi:10.1016/j.imbio.2008.11.007

11. Garlanda C, Di Ceglie I, Jaillon S. IL-1 family cytokines in inflammation and immunity. Cell Mol Immunol. 2025 Oct 14;22(11):1345–62. doi:10.1038/s41423-025-01358-8

12. Monteleone M, Stanley AC, Chen KW, Brown DL, Bezbradica JS, Von Pein JB, et al. Interleukin-1β Maturation Triggers Its Relocation to the Plasma Membrane for Gasdermin-D-Dependent and –Independent Secretion. Cell Reports. 2018 Aug;24(6):1425–33. doi:10.1016/j.celrep.2018.07.027

13. Brough D, Rothwell NJ. Caspase-1-dependent processing of pro-interleukin-1β is cytosolic and precedes cell death. Journal of Cell Science. 2007 Mar 1;120(5):772–81. doi:10.1242/jcs.03377

14. Groß O, Yazdi AS, Thomas CJ, Masin M, Heinz LX, Guarda G, et al. Inflammasome Activators Induce Interleukin-1α Secretion via Distinct Pathways with Differential Requirement for the Protease Function of Caspase-1. Immunity. 2012 Mar;36(3):388–400. doi:10.1016/j.immuni.2012.01.018

15. Van Opdenbosch N, Lamkanfi M. Caspases in Cell Death, Inflammation, and Disease. Immunity. 2019 Jun 18;50(6):1352–64. doi:10.1016/j.immuni.2019.05.020 PubMed PMID: 31216460; PubMed Central PMCID: PMC6611727.

16. Néel E, Chiritoiu-Butnaru M, Fargues W, Denus M, Colladant M, Filaquier A, et al. The endolysosomal system in conventional and unconventional protein secretion. Journal of Cell Biology. 2024 Sep 2;223(9):e202404152. doi:10.1083/jcb.202404152

17. Malhotra V. Unconventional protein secretion: an evolving mechanism. EMBO J. 2013 Jun 12;32(12):1660–4. doi:10.1038/emboj.2013.104

18. Jacopo M. Unconventional protein secretion (UPS): role in important diseases. Mol Biomed. 2023 Jan 9;4(1):2. doi:10.1186/s43556-022-00113-z

19. Broz P. Unconventional protein secretion by gasdermin pores. Seminars in Immunology. 2023 Sep;69:101811. doi:10.1016/j.smim.2023.101811

20. Sborgi L, Rühl S, Mulvihill E, Pipercevic J, Heilig R, Stahlberg H, et al. GSDMD membrane pore formation constitutes the mechanism of pyroptotic cell death. The EMBO Journal. 2016 Aug 15;35(16):1766–78. doi:10.15252/embj.201694696

21. Broz P, Pelegrín P, Shao F. The gasdermins, a protein family executing cell death and inflammation. Nat Rev Immunol. 2020 Mar;20(3):143–57. doi:10.1038/s41577-019-0228-2

22. Broz P. Caspase target drives pyroptosis. Nature. 2015 Oct 29;526(7575):642–3. doi:10.1038/nature15632

23. Chiritoiu M, Brouwers N, Turacchio G, Pirozzi M, Malhotra V. GRASP55 and UPR Control Interleukin-1β Aggregation and Secretion. Dev Cell. 2019 Apr 8;49(1):145–155.e4. doi:10.1016/j.devcel.2019.02.011 PubMed PMID: 30880003.

24. Arkusz J, Stepnik M, Trzaska D, Dastych J, Rydzyński K. Assessment of usefulness of J774A.1 macrophages for the assay of IL-1β promoter activity. Toxicology in Vitro. 2006 Feb;20(1):109–16. doi:10.1016/j.tiv.2005.06.044

25. Oh-hashi K, Furuta E, Fujimura K, Hirata Y. Application of a novel HiBiT peptide tag for monitoring ATF4 protein expression in Neuro2a cells. Biochemistry and Biophysics Reports. 2017 Dec;12:40–5. doi:10.1016/j.bbrep.2017.08.002

26. Schwinn MK, Machleidt T, Zimmerman K, Eggers CT, Dixon AS, Hurst R, et al. CRISPR-Mediated Tagging of Endogenous Proteins with a Luminescent Peptide. ACS Chem Biol. 2018 Feb 16;13(2):467–74. doi:10.1021/acschembio.7b00549

27. Edye ME, Brough D, Allan SM. Acid-dependent Interleukin-1 (IL-1) Cleavage Limits Available Pro-IL-1β for Caspase-1 Cleavage. Journal of Biological Chemistry. 2015 Oct;290(42):25374–81. doi:10.1074/jbc.M115.667162

28. Vijayaraj SL, Feltham R, Rashidi M, Frank D, Liu Z, Simpson DS, et al. The ubiquitylation of IL-1β limits its cleavage by caspase-1 and targets it for proteasomal degradation. Nat Commun. 2021 May 11;12(1):2713. doi:10.1038/s41467-021-22979-3

29. Chaffey LE, Roberti A, Bowman A, O’Brien CJo, Som L, Purvis GSd, et al. Drug repurposing screen identifies novel anti-inflammatory activity of sunitinib in macrophages. European Journal of Pharmacology. 2024 Apr;969:176437. doi:10.1016/j.ejphar.2024.176437

30. Xu J, Pickard JM, Núñez G. FDA-approved disulfiram inhibits the NLRP3 inflammasome by regulating NLRP3 palmitoylation. Cell Reports. 2024 Aug;43(8):114609. doi:10.1016/j.celrep.2024.114609

31. King, Jr JG, Khalili K. Inhibition of human brain tumor cell growth by the anti-inflammatory drug, flurbiprofen. Oncogene. 2001 Oct 18;20(47):6864–70. doi:10.1038/sj.onc.1204907

32. Ortonne J -P., Esposito M, Chimenti S, Kapińska-Mrowiecka M, Grodzińska A, Naldi L, et al. Betamethasone valerate dressing is non-inferior to calcipotriol–betamethasone dipropionate ointment in the treatment of patients with mild-to-moderate chronic plaque psoriasis: results of a randomized assessor-blinded multicentre trial. Acad Dermatol Venereol. 2014 Sep;28(9):1226–34. doi:10.1111/jdv.12270

33. Denus M, Filaquier A, Fargues W, Néel E, Stewart SE, Colladant M, et al. A Sensitive and Versatile Cell-Based Assay Combines Luminescence and Trapping Approaches to Monitor Unconventional Protein Secretion. Traffic. 2025 Apr;26(4–6):e70009. doi:10.1111/tra.70009

34. Néel E, Denus M, Fargues W, Gal C, Enjolras C, Boulanger A, et al. Protocols for Monitoring Unconventional Protein Secretion Using Luminescence and Trapping Approaches. Current Protocols. 2026 Feb;6(2):e70326. doi:10.1002/cpz1.70326

35. Guay DRP. Tolcapone, a Selective Catechol-*O* –Methyltransferase Inhibitor for Treatment of Parkinson’s Disease. Pharmacotherapy. 1999 Jan 2;19(1):6–20. doi:10.1592/phco.19.1.6.30516

36. Yan Y, Jiang W, Liu L, Wang X, Ding C, Tian Z, et al. Dopamine Controls Systemic Inflammation through Inhibition of NLRP3 Inflammasome. Cell. 2015 Jan;160(1–2):62–73. doi:10.1016/j.cell.2014.11.047

37. Mariathasan S, Weiss DS, Newton K, McBride J, O’Rourke K, Roose-Girma M, et al. Cryopyrin activates the inflammasome in response to toxins and ATP. Nature. 2006 Mar;440(7081):228–32. doi:10.1038/nature04515

38. Sutterwala FS, Ogura Y, Szczepanik M, Lara-Tejero M, Lichtenberger GS, Grant EP, et al. Critical Role for NALP3/CIAS1/Cryopyrin in Innate and Adaptive Immunity through Its Regulation of Caspase-1. Immunity. 2006 Mar;24(3):317–27. doi:10.1016/j.immuni.2006.02.004

39. Saccani S, Polentarutti N, Penton-Rol G, Sims JE, Mantovani A. DIVERGENT EFFECTS OF LPS ON EXPRESSION OF IL-1 RECEPTOR FAMILY MEMBERS IN MONONUCLEAR PHAGOCYTES IN VITRO AND IN VIVO. Cytokine. 1998 Oct;10(10):773–80. doi:10.1006/cyto.1998.0359

40. Duncan LM, Meegan LS, Unanue ER. IL-1 gene expression in lymphoid tissues. The Journal of Immunology. 1991 Jan 15;146(2):565–71. doi:10.4049/jimmunol.146.2.565

41. Liu T, Wang Q, Du Z, Yin L, Li J, Meng X, et al. The trigger for pancreatic disease: NLRP3 inflammasome. Cell Death Discov. 2023 Jul 14;9(1):246. doi:10.1038/s41420-023-01550-7

42. Forsberg M, Lehtonen M, Heikkinen M, Savolainen J, Järvinen T, Männistö PT. Pharmacokinetics and Pharmacodynamics of Entacapone and Tolcapone after Acute and Repeated Administration: A Comparative Study in the Rat. The Journal of Pharmacology and Experimental Therapeutics. 2003 Feb;304(2):498–506. doi:10.1124/jpet.102.042846

43. Smith KS, Smith PL, Heady TN, Trugman JM, Harman WD, Macdonald TL. In Vitro Metabolism of Tolcapone to Reactive Intermediates: Relevance to Tolcapone Liver Toxicity. Chem Res Toxicol. 2003 Feb 1;16(2):123–8. doi:10.1021/tx025569n

44. Landy E, Carol H, Ring A, Canna S. Biological and clinical roles of IL-18 in inflammatory diseases. Nat Rev Rheumatol. 2024 Jan;20(1):33–47. doi:10.1038/s41584-023-01053-w

45. Di Paolo NC, Shayakhmetov DM. Interleukin 1α and the inflammatory process. Nat Immunol. 2016 Aug;17(8):906–13. doi:10.1038/ni.3503

46. Tapia VS, Daniels MJD, Palazón-Riquelme P, Dewhurst M, Luheshi NM, Rivers-Auty J, et al. The three cytokines IL-1β, IL-18, and IL-1α share related but distinct secretory routes. Journal of Biological Chemistry. 2019 May;294(21):8325–35. doi:10.1074/jbc.RA119.008009

47. Puren AJ, Fantuzzi G, Dinarello CA. Gene expression, synthesis, and secretion of interleukin 18 and interleukin 1β are differentially regulated in human blood mononuclear cells and mouse spleen cells. Proc Natl Acad Sci USA. 1999 Mar 2;96(5):2256–61. doi:10.1073/pnas.96.5.2256

48. Pirhonen J, Sareneva T, Kurimoto M, Julkunen I, Matikainen S. Virus Infection Activates IL-1β and IL-18 Production in Human Macrophages by a Caspase-1-Dependent Pathway. The Journal of Immunology. 1999 Jun 15;162(12):7322–9. doi:10.4049/jimmunol.162.12.7322

49. Verweyen E, Holzinger D, Weinhage T, Hinze C, Wittkowski H, Pickkers P, et al. Synergistic Signaling of TLR and IFNα/β Facilitates Escape of IL-18 Expression from Endotoxin Tolerance. American Journal of Respiratory and Critical Care Medicine. 2020 Mar 1;201(5):526–39. doi:10.1164/rccm.201903-0659OC

50. Fontana P, Du G, Zhang Y, Zhang H, Vora SM, Hu JJ, et al. Small-molecule GSDMD agonism in tumors stimulates antitumor immunity without toxicity. Cell. 2024 Oct;187(22):6165–6181.e22. doi:10.1016/j.cell.2024.08.007

51. Zhang Z, Zhang Y, Xia S, Kong Q, Li S, Liu X, et al. Gasdermin E suppresses tumour growth by activating anti-tumour immunity. Nature. 2020 Mar 19;579(7799):415–20. doi:10.1038/s41586-020-2071-9

52. Neel DV, Basu H, Gunner G, Bergstresser MD, Giadone RM, Chung H, et al. Gasdermin-E mediates mitochondrial damage in axons and neurodegeneration. Neuron. 2023 Apr;111(8):1222–1240.e9. doi:10.1016/j.neuron.2023.02.019

53. Churchill MJ, Mitchell PS, Rauch I. Epithelial Pyroptosis in Host Defense. J Mol Biol. 2022 Feb 28;434(4):167278. doi:10.1016/j.jmb.2021.167278 PubMed PMID: 34627788; PubMed Central PMCID: PMC10010195.

54. Coll RC, Hill JR, Day CJ, Zamoshnikova A, Boucher D, Massey NL, et al. MCC950 directly targets the NLRP3 ATP-hydrolysis motif for inflammasome inhibition. Nat Chem Biol. 2019 Jun;15(6):556–9. doi:10.1038/s41589-019-0277-7

55. Olanow CW, Watkins PB. Tolcapone: An Efficacy and Safety Review (2007). Clinical Neuropharmacology. 2007 Sep;30(5):287–94. doi:10.1097/wnf.0b013e318038d2b6

56. Olanow CW. Tolcapone and Hepatotoxic Effects. Arch Neurol. 2000 Feb 1;57(2):263. doi:10.1001/archneur.57.2.263

57. Vieira-Coelho MA, Soares-da-Silva P. Effects of tolcapone upon soluble and membrane-bound brain and liver catechol-O-methyltransferase. Brain Research. 1999 Mar;821(1):69–78. doi:10.1016/S0006-8993(99)01063-X

58. Müller T. Entacapone/Tolcapone/Opicapone for Treating Parkinson’s Disease. In: Riederer P, Laux G, Nagatsu T, Le W, Riederer C, editors. NeuroPsychopharmacotherapy [Internet]. Cham: Springer International Publishing; 2022 [cited 2026 Mar 6]. p. 3309–26. Available from: https://link.springer.com/10.1007/978-3-030-62059-2_230 doi:10.1007/978-3-030-62059-2_230

59. Rathkey JK, Zhao J, Liu Z, Chen Y, Yang J, Kondolf HC, et al. Chemical disruption of the pyroptotic pore-forming protein gasdermin D inhibits inflammatory cell death and sepsis. Sci Immunol. 2018 Aug 3;3(26):eaat2738. doi:10.1126/sciimmunol.aat2738

60. Hu Y, Li H, Zhang X, Song Y, Liu J, Pu J, et al. Identification of two repurposed drugs targeting GSDMD oligomerization interface I to block pyroptosis. Cell Chemical Biology. 2024 Dec;31(12):2024–2038.e7. doi:10.1016/j.chembiol.2024.10.002

61. Dhani S, Zhao Y, Zhivotovsky B. A long way to go: caspase inhibitors in clinical use. Cell Death Dis. 2021 Oct 15;12(10):949. doi:10.1038/s41419-021-04240-3

62. Lopez-Castejon G, Theaker J, Pelegrin P, Clifton AD, Braddock M, Surprenant A. P2X7 Receptor-Mediated Release of Cathepsins from Macrophages Is a Cytokine-Independent Mechanism Potentially Involved in Joint Diseases. The Journal of Immunology. 2010 Aug 15;185(4):2611–9. doi:10.4049/jimmunol.1000436

